# Phencyclidine discoordinates hippocampal network activity but not place fields

**DOI:** 10.1101/200162

**Authors:** Hsin-Yi Kao, Dino Dvořák, EunHye Park, Jana Kenney, Eduard Kelemen, André A Fenton

## Abstract

We used the psychotomimetic phencyclidine (PCP) to investigate the relationships between cognitive behavior, coordinated neural network function and information processing within the hippocampus place cell system. We report in rats that PCP (5mg/kg i.p.) impairs a well-learned hippocampus-dependent place avoidance behavior in rats that requires cognitive control, even when PCP is injected directly into dorsal hippocampus. PCP increases 60-100 Hz medium gamma oscillations in hippocampus CA1 and these increases correlate with the cognitive impairment caused by systemic PCP administration. PCP discoordinates theta-modulated medium and slow gamma oscillations in CA1 local field potentials (LFP) such that medium gamma oscillations become more theta-organized than slow gamma oscillations. CA1 place cell firing fields are preserved under PCP but the drug discoordinates the sub-second temporal organization of discharge amongst place cells. This discoordination causes place cell ensemble representations of a familiar space to cease resembling pre-PCP representations, despite preserved place fields. These findings point to the cognitive impairments caused by PCP arising from neural discoordination. PCP disrupts the timing of discharge with respect to the sub-second timescales of theta and gamma oscillations in the LFP. Because these oscillations arise from local inhibitory synaptic activity, these findings point to excitation-inhibition discoordination as the root of PCP-induced cognitive impairment.

**SIGNIFICANCE STATEMENT:** Hippocampal neural discharge is temporally coordinated on timescales of theta and gamma oscillations in the local field potential, and the discharge of a subset of pyramidal neurons called “place cells” is spatially organized such that discharge is restricted to locations called a cell’s “place field.” Because this temporal coordination and spatial discharge organization is thought to represent spatial knowledge, we used the psychotomimetic phencyclidine (PCP) to disrupt cognitive behavior and assess the importance of neural coordination and place fields for spatial cognition. PCP impaired the judicious use of spatial information and discoordinated hippocampal discharge, without disrupting firing fields. These findings dissociate place fields from spatial cognitive behavior and suggest that hippocampus discharge coordination is crucial to spatial cognition.

## INTRODUCTION

Place cells are hippocampus principal cells that discharge in ‘place fields’ that map discharge to locations, making place cell studies *de facto* investigations of what information the hippocampus represents and how that information is represented (Friston and Buzsaki, 2016). Place field stability, while not absolute (Fenton et al., 2010; Ziv et al., 2013), is taken as crucial evidence that hippocampus discharge represents spatial memory and acquired knowledge (Muller et al., 1987; Lever et al., 2002; Kentros et al., 2004; Wills et al., 2005).

The discharge of multiple place cells is temporally coordinated such that the likelihood that two cells will discharge in sub-second temporal proximity is characteristic for a cell pair (Dragoi and Buzsaki, 2006; Kelemen and Fenton, 2012). These temporal discharge patterns may themselves be crucial for place field firing and may in fact represent the information that animals use for spatial cognition (Buzsaki, 2010). Accordingly, manipulations that sufficiently disturb the temporal discharge relationships amongst cells, what is called ‘neural coordination,’ should also disturb place fields, and *vice versa*, and such manipulations should also disrupt spatial cognitive behavior. However, the ‘discoordination’ hypothesis predicts that aberrant neural coordination and impaired cognition can associate with intact place fields. According to it, cognitive deficits such as schizophrenia-related impaired cognitive control, arise because processing multiple streams of information is corrupted by discoordination of neural activity, in which cells that normally discharge together and cells that normally do not discharge together fail to maintain their appropriate timing relationships, despite maintaining their individual response properties (Tononi and Edelman, 2000; Phillips and Silverstein, 2003; Uhlhaas and Singer, 2006; Lee et al., 2012; Lee et al., 2014; O’Reilly et al., 2014; Fenton, 2015).

The discoordination hypothesis derives from cell assembly (Hebb, 1949) and other ensemble hypotheses for how information is represented in the brain, which seem valid in hippocampus (Harris et al., 2003; Itskov et al., 2008; Fenton et al., 2010; Kelemen and Fenton, 2010; Park et al., 2011). Ensemble hypotheses assert that streams of information are represented by the coordinated temporal spiking relationships amongst cells in distributed representations. It follows that temporal discoordination will derange ensemble representations and the judicious use of information.

Here we use a standard place cell paradigm with the psychotomimetic phencyclidine (PCP) to disturb cognition so as to evaluate the discoordination hypothesis by investigating the relationships between place fields, neural coordination, and spatial cognition (Fig. 1). PCP causes thought disorder in healthy people and exacerbates symptoms in schizophrenia patients (Cohen et al., 1962; Itil et al., 1967; Abi-Saab et al., 1998), but despite a clear pharmacological mechanism of action, how PCP causes cognitive abnormalities is unknown. PCP is a potent uncompetitive antagonist of the glutamatergic N-methyl-D-aspartate receptor (NMDAR) with dopamine-mimetic effects at D2 as well as other receptors (Lodge and Anis, 1982; Anis et al., 1983). Although blocking NMDARs reduces excitation, it is unclear whether PCP’s consequences arise from hypo- or hyperglutaminergic effects (Adams and Moghaddam, 1998; Seeman et al., 2005; Sershen et al., 2008). Indeed, there is substantial evidence of network activation by PCP and other psychotomimetic NMDAR antagonists, dizocilpine (MK801) and ketamine (Benardo, 1995; Grunze et al., 1996). These drugs increase activity of prefrontal (Moghaddam et al., 1997) and cingulate cortical areas (Breier et al., 1997; Vollenweider et al., 1997) in human neuroimaging studies. In rats, MK801 decreases interneuron firing and increases pyramidal cell firing in prefrontal areas (Holcomb et al., 2005; Rowland et al., 2005) and disrupts the coupling of spiking to local gamma oscillations (Homayoun and Moghaddam, 2007) consistent with disinhibition effects. We stress that we use PCP, neither to model schizophrenia nor to investigate a particular molecular component of hippocampus function, rather we use PCP as a potent means of disrupting hippocampus-dependent cognitive function and ask whether the drug also disrupts the single cell (firing fields) and ensemble (diverse neural coordination phenomena) discharge properties of the hippocampus.

**Figure 1.**
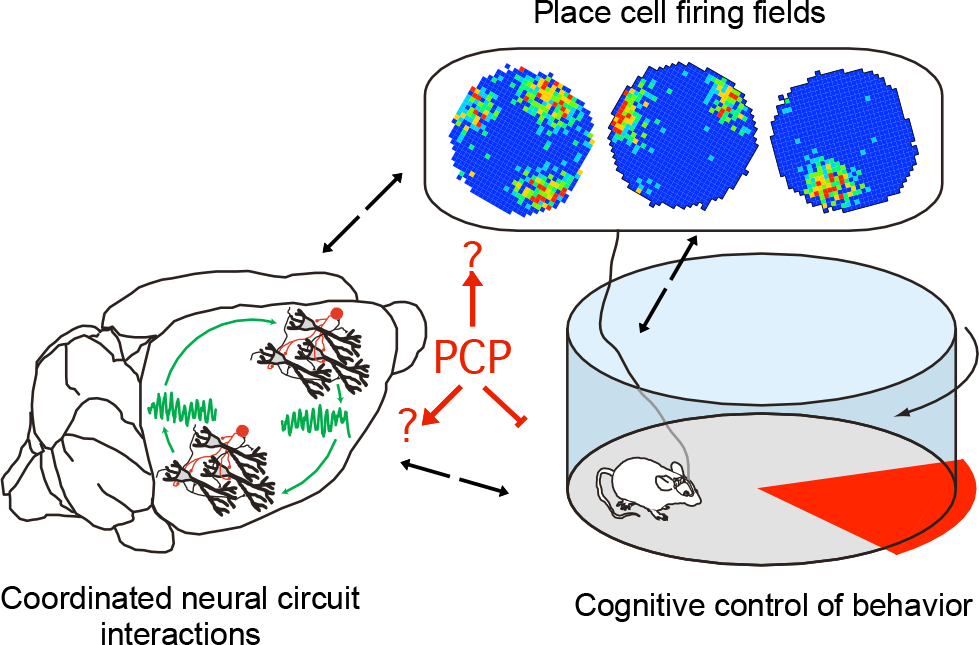
Three levels of neurobiological function are assessed to examine widely-held assumptions of their interrelatedness. Diverse prior work has assumed that neural coordination within the hippocampus circuit is crucial for generating place cell firing fields and the effective use of spatial information; firing fields are thought to be crucial for the encoding and effective use of spatial information. The neurobiological mechanisms at each level of analysis are assumed to feedback to impact the other levels. This perspective, often implicit in the field, assumes that a manipulation to induce cognitive dysfunction manifest as impaired use of spatial information, will also disrupt neural coordination and place cell firing fields. PCP is well documented to impair cognition, but these assumed impacts of PCP have not been rigorously evaluated. Here they are evaluated together using the active place avoidance paradigm to assess hippocampus-dependent information processing, and the hippocampus place cell paradigm to assess cognitive representations and coordinated hippocampus function.

## MATERIALS AND METHODS

All behavioral and electrophysiological methods have been published elsewhere, and are only briefly described here.

### Animals

All animals were maintained on a 12:12 light-dark cycle. Adult male Long-Evans rats were 2-3 months during the experiments. All procedures conformed to institutional and NIH guidelines for the ethical treatment of vertebrate animals and were approved by the SUNY, Downstate and NYU Institutional Animal Care and Use Committees as well as the ethics committee of the Institute of Physiology, Czech Academy of Sciences in accord with the directive of the European Community Council (86/609/ECC).

### Drugs

Phencyclidine (PCP, 1-(1-Phenylcyclohexyl)piperidine hydrochloride, pH 7.4; Sigma, St. Louis, MO) solutions were made fresh in sterile saline before use.

### Active place avoidance task - Setup

An individual animal was placed on a metal 82-cm diameter disk-shaped arena see Fig. 2A). The arena was elevated 76 cm above the floor and centered in a 3×4 m^2^ room surrounded by opaque curtains and various items. The animal was constrained to remain on the disk by a 40-cm-high transparent wall that allows the animal to see the surrounding room through the wall. The position of the animal was tracked every 17 ms from an overhead camera using digital video spot tracking software (Tracker, Bio-Signal Group Corp., Acton, MA, USA).

**Figure 2.**
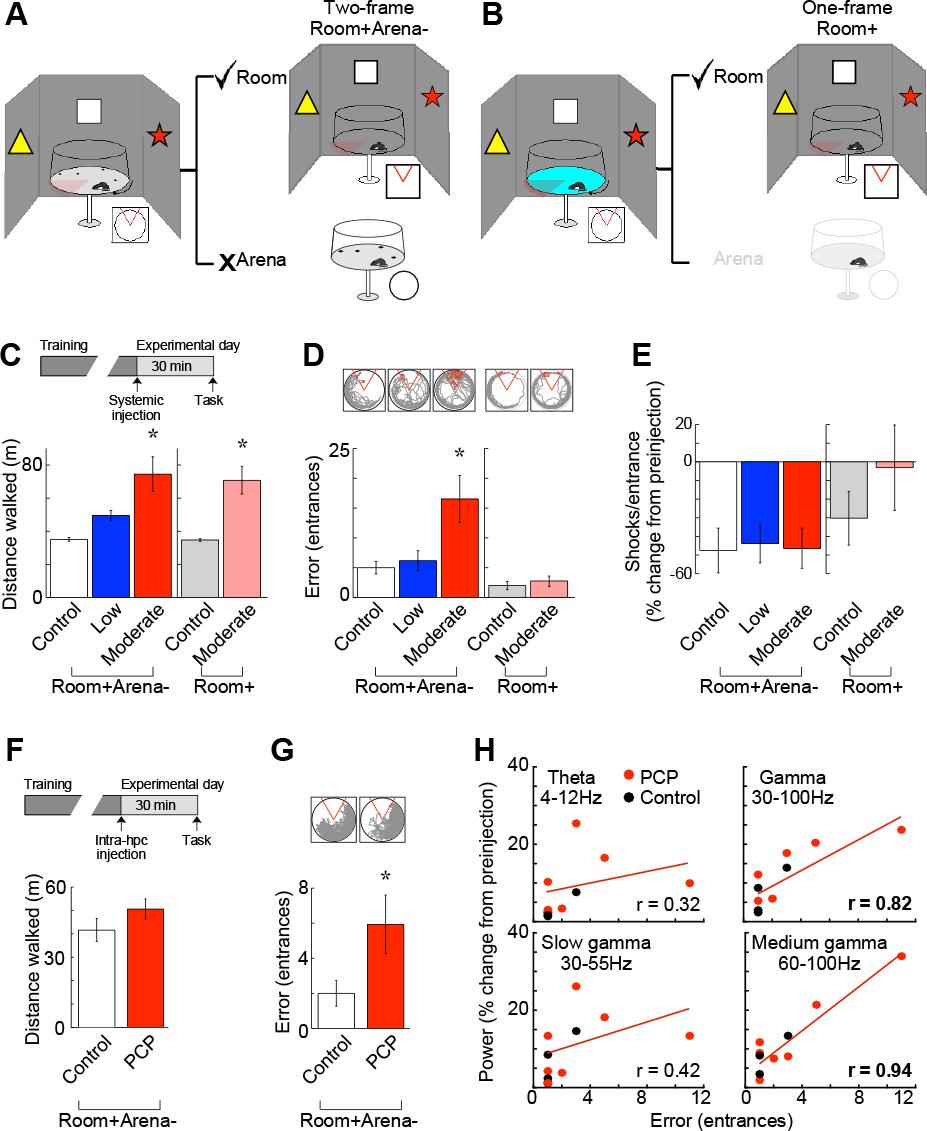
PCP impairs cognitive control. Schematics of the active place avoidance tasks on a continuously rotating arena that creates two distinct frames of spatial information, one defined by stationary room cues and another defined by rotating arena cues. (A) In the two-frame, Room+Arena- task variant, the rat must avoid a stationary shock zone (red) by using (✓) the stream of relevant ‘Room’ information from the stationary frame, while ignoring (✗) the stream of irrelevant ‘Arena’ information from the rotating frame. Using one class of information while ignoring another requires cognitive control. (B) In the one-frame, Room+ task variant, the irrelevant ‘Arena’ information can be minimized by shallow water to attenuate the arena cues for place localization, and reduce the demand for cognitive control. (C) Inset: The experimental design. Rats were trained for several days on one task variant. After optimal performance, the rats received systemic administration of either a control or PCP dose 30 minutes before testing. The moderate dose of PCP causes hyperlocomotion, measured by the distance that rat walked during both task variants. (D) Inset: Representative paths from a single rat; red indicates the shock zone and shocks. The moderate dose of PCP impairs familiar two-frame (n = 10), but not one-frame place avoidance (n = 4), measured by the number of errors, entrances to the shock zone. (E) PCP did not change responses to shock, measured as the change from the initial experience of shock prior to PCP treatment. The number of shocks/entrance estimates the rat’s willingness to escape shock. (F) Infusing PCP into both hippocampi (inset: Experimental design n = 11). Intrahippocampal PCP does not cause hyperlocomotion, but (G) is sufficient to impair familiar two-frame Room+Arena– place avoidance (Inset: representative paths from a single rat). (H) The Room+Arena– avoidance deficit following intrahippocampal infusion was related to the magnitude of the change in medium gamma oscillation amplitude in the hippocampal LFP, but not the changes in theta or slow gamma oscillation amplitudes (n = 8). Correlations and regression lines are for fits only to the PCP data; bold indicates significance. Error bars and shading: ± S.E.M.; Control, low, moderate: 0, 3, 5mg/kg PCP doses. * indicates p < 0.05 relative to all groups. Bold type indicates significant correlations.

Both a two-frame Room+Arena- task variant (Fig. 2A) and a one-frame Room+ task variant (Fig. 2B) were used. In both tasks the arena rotated at 1 rpm and an unmarked 60° shock zone was fixed in stationary room coordinates. To avoid shock the animal had to use the relevant stationary distal room cues to localize itself and the positions of shock. Because the arena was rotating the animal also had to ignore the irrelevant information from the rotating spatial frame. In the Room+Arena- task variant the irrelevant cues were scents and fecal boli that were stable on the arena surface, but in the Room+ variant their salience was reduced by shallow water (~2 cm saline) on the arena surface (Wesierska et al., 2005). Thus in the Room+Arena– task variant the demand for cognitive control to ignore the salient irrelevant arena cues was high but the demand was low in the Room+ task. All other cognitive and motivational aspects of the two tasks were similar. Avoidance was reinforced by a constant current foot-shock (0.3 mA, 60 Hz, 500 ms). The shock was delivered across the rat on the grounded arena surface via a low impedance stainless steel shock electrode that was implanted subcutaneously between the rat’s shoulders so the major voltage drop was across the rat’s paws. Under control of software, shock was delivered when the animal was in the shock zone for 500 msec. The shock was repeated every 1.5 seconds until the animal left the shock zone. The spatial resolution of the position measurements was ~ 3 mm.

### Active place avoidance task - Behavioral Protocol

The goal was to test the effect of PCP on active place avoidance after the task had been learned and the memories established. Rats were first habituated to the environment during a 10-min trial on the stationary arena with no shock. The shock and arena rotation were then turned on and rats were given five 10-min training trials with 10-min inter-trial interval. The 5 trials/day training protocol continued on subsequent days until the rat’s performance reached a criterion number of shock zone entries, which was ≤ 2 in two consecutive trials. The day after reaching criterion, each rat received a dose of PCP by either systemic injection or intrahippocampal infusion 30 min before testing in conditions that were identical to the training. The rats had a wash-out period of at least 48 h between different doses of PCP.

The time series of the animal’s positions was analyzed. The distance the animals walked was used to quantify locomotion. This measure increased after systemic PCP administration because the drug causes hyperlocomotion. The number of entrances into the shock zone was counted to estimate place avoidance. This measure decreases as the animals learn to avoid shock and can be interpreted as an index of the ability to perform the conditioned avoidance and judicious retrieval of the place avoidance memory in both task variants (Wesierska et al., 2005; Lee et al., 2012). To estimate the animal’s willingness to escape shock under PCP, we measured the number of shocks that were received each time the shock zone was entered. Since this measure depends on the subject’s prior experience with shock, we computed the difference in the shocks/entrance during the PCP trial compared to the initial trial before any experience with shock and PCP. In addition to comparing the measure of place avoidance in the high and low cognitive control task variants, this analysis of shocks/entrance served as additional evidence for evaluating whether or not PCP impaired place avoidance *per se*, or merely the willingness to avoid shock itself.

### Intrahippocampal infusion - Surgical preparation and infusion protocol

Rats were prepared for intrahippocampal infusions according to published methods (Wesierska et al., 2005; Pastalkova et al., 2006). Briefly, under Nembutal anesthesia (50 mg/kg) a pair of 22-Ga guide cannulae were stereotaxically implanted and secured by dental cement above the dorsal hippocampal injection targets (AP: -4.0 mm, ML: ±2.5 mm, DV: -2.0 mm below the dura). Behavioral training began at least 1 week after surgery. PCP solution was infused at the hippocampal targets by pressure injection using a Hamilton syringe connected to a 30-Ga needle by Tygon tubing. The rat was restrained by hand for infusions. One μl of saline (0.9 %) or PCP (6, 8, and 10 μg/μl) was infused into each site during 2 min. The needle was withdrawn 2 min after infusion ended. The rat was then returned to the home cage and observed for 30 min before testing.

### *In vivo* electrophysiology - Acute recordings under anesthesia

The *in vivo* methods to record single unit discharge from urethane-anesthetized rats have been described in prior work (Olypher et al., 2006). Briefly, Rats were anesthetized with urethane (1.2g/kg i.p.) then mounted in a stereotaxic frame with precision micromanipulators (TSE Systems GmbH, Bad Homburg, Germany). The scalp was resected and two crainotomies were performed, one at AP -3.8 mm, ML 2.5 mm relative to bregma to provide electrode access to the dorsal hippocampus and the other at AP 3.0 mm, ML 0.5-0.7 mm to provide access to the medial prefrontal cortex. The dura was cut and tetrode-configured electrodes made from 4 twisted 25-μm nichrome wires (impedance 50-200 kΩ), were lowered to the recording targets areas using the stereotaxic micromanipulators, guided by electrophysiological landmarks in the local field potentials and single unit activity. The rest of the recording setup was the same as for the chronic recordings (see below). Data collection began once stable single unit ensemble activity could be isolated. A baseline recording was made for at least 30 minutes and then the rat received an i.p. injection of the 5 mg/kg PCP or vehicle solution. Recordings continued at least one hour after the injection.

### *In vivo* electrophysiology - Surgical preparation for chronic recordings

The *in vivo* electrophysiological methods have been previously described (Kelemen and Fenton, 2010). Briefly, tetrodes were made from 4 twisted 25-μm nichrome wires (impedance 50-200 kΩ). Eight tetrodes were assembled in a custom microdrive. The microdrive was stereotaxically implanted in rats under Nembutal anesthesia (50 mg/kg) and fixed to the skull with bone screws and dental cement. The tetrodes were aimed at a location above the left dorsal hippocampus (AP: -4.0 mm, ML: -2.5 mm, DV: -1.9 mm below the brain surface). At least 1 week after surgery, the tetrodes were gradually moved toward the hippocampus until action potentials from CA1 place cells could be recorded.

### *In vivo* electrophysiology - Recording setup

Extracellular single-unit activity was recorded from the rats using a cable that made electrical contact with the recording electrodes via a Millmax connector integrated into the electrode assembly. Extracellular, 2-ms tetrode action potential waveforms were buffered by a custom preamplifier, band-pass filtered (low-cut frequency 300 or 360 Hz, hi-cut frequency 5-10 kHz), amplified (5,000-10,000 times), and digitized (32 or 48 kHz). The digital signals were stored in real-time using either custom software (AcX, A.A. Fenton) or a commercial system (dacqUSB, Axona Ltd., St. Albans U.K). Local field potentials were also acquired from an independently placed tetrode or 75-μm nichrome electrode. Single unit signals were referenced to this local electrode and LFP signals were referenced to an electrode that was targeted to the cerebellar white matter. The LFP signals were band-pass filtered (0.1-300 Hz or 0.1-360 Hz) and digitized at 2 kHz.

For recordings from awake behaving rats, the electrophysiological data were recorded in parallel and synchronized with the rat’s position on the arena.

Place cell recordings were made as the rats foraged on the arena for randomly scattered sugar pellets (Bio-serve, Frenchtown, NJ). Once an ensemble of action potential spike trains from 8 or more place cells could be recorded, an experiment began. The ensemble was recorded for at least 30 min as the rats foraged continuously in the pre-injection session. The rat was returned to the home cage and provided with water for 10 min, then received an i.p. injection of the control 0 mg/kg, low 3 mg/kg, or moderate 5mg/kg dose of PCP before being returned to the arena. Postinjection recordings lasted 60 min. All data were analyzed off-line. The data recorded in the 30– min pre-injection and the first 30-min post-injection intervals were analyzed for statistical comparisons.

### *In vivo* electrophysiology - Local field potential recording during foraging for scattered food

Power spectra were computed for the 30-min pre-injection and the first 30-min post-injection recordings using Matlab’s Pwelch function. The cross-frequency coupling was estimated between the phase of slow frequency oscillations (2-12 Hz) and the amplitude of faster oscillations (25-250 Hz). We used the algorithm introduced by Canolty et al. (Canolty et al., 2006) with filters designed by Matlab’s Filter Design Toolbox (Dvorak and Fenton, 2014).

Phase-frequency discharge probability plots (Fig. 7) were computed by convolving the LFP signal with a group of complex Morlet wavelets in the logarithmic range between 2 and 250 Hz. Instantaneous phase of the band-specific LFP signals was obtained from the complex time series. The oscillation phase discharge probability was computed independently for each frequency band. The frequency-specific phase distribution of spiking was normalized by dividing the discharge distribution by the total number of action potentials in a given recording.

### *In vivo* electrophysiology - Local field potential recording during place avoidance

Electrodes were made from 6 twisted 75-μm nichrome wires (impedance 50-200 kΩ) cut at an angle (1 mm between 1^st^ and 6^th^ wires) as in prior work (Radwan et al., 2016). The electrode was stereotaxically implanted in rats under Nembutal anesthesia (50 mg/kg) and fixed to the skull with bone screws and dental cement when the cannulae for intrahippocampal infusion were implanted (Kelemen and Fenton, 2010). The tip of the electrode was aimed in the left dorsal hippocampus CA1 (AP: -4.0 mm, ML: -2.5 mm, DV: -3.0 mm below the dura). The rats were tested in the behavioral task at least one week after surgery.

Power spectra were computed for the recordings of the last training trial and the first trial after PCP treatment using Matlab’s Pwelch function.

### *In vivo* electrophysiology - Single unit identification

The action potentials emitted by different cells were discriminated off-line using custom waveform parameter clustering software (Wclust, A.A. Fenton). Each action potential waveform was characterized by parameters that included the positive and negative peak voltages on each tetrode wire, the voltage at user-selected times relative to the spike onset, the waveform energy, principal components, and others. Single units were classified as those waveforms that formed distinctive clusters in the waveform parameter space according to objective criteria (Neymotin et al., 2011).

### Electrophysiological data analysis - Classification of cell types

Single units were classified as complex-spike or theta cells according to published criteria (Ranck, 1973; Fenton et al., 2008). Complex-spike cells appear to be pyramidal cells, whereas theta cells are likely local interneurons (Fox and Ranck, 1975). Pyramidal cells were judged to have long-duration waveforms (> 250 μs), low discharge rate (< 2 AP/s) and a tendency to fire in bursts (peak inter-spike interval < 10 ms). Interneurons had short-duration waveforms (< 250 μs), high discharge rate (> 2 AP/s), and did not tend to fire in bursts.

### Electrophysiological data analysis - Characterization of place cells

Cell-specific spatial firing rate maps were created for each cell by calculating the total number of spikes observed in each 2.5×2.5 cm^2^ location divided by the total time the rat was observed in each corresponding location (examples are given in Fig. 1). Standard measures of spatial firing quality were then computed from each cell’s firing rate map (Fenton et al., 2008). We report three of these measures: the overall firing rate is the total spikes a cell discharged divided by the total recording time; the spatial coherence describes the local smoothness of the firing rate distribution (Muller and Kubie, 1989); the spatial information content describes the reduction in uncertainty about the rat’s position given a particular firing rate (Skaggs et al., 1993).

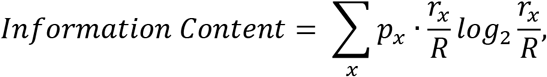

where r_x_ is the mean rate in pixel x; R is the overall mean rate; p_x_ is the probability of finding the rat in pixel x and p_x_ = t_x_/T where t_x_ is the time spent in pixel x and T is the total recording time.

Single units were classified as place cells if the overall discharge rate was (0.1-2 AP/s), the spatial coherence (z score) was > 0.4, and the information content was > 0.4 bits/spike. The spatial similarity was also computed. It describes how similar two firing rate maps are. It is computed as Pearson’s correlation for the firing rates in corresponding pixels of the two maps.

### Electrophysiological data analysis - Standardization of positional sampling

PCP is well known to cause hyperactivity and stereotypy in the movements of rodents, which could influence the quantitative measures of spatial firing quality (Muller et al., 1987). Accordingly, we standardized each place cell data time series to minimize positional sampling differences between recordings under the different doses of PCP. The goal was to make the standardized maps intermediate between a pre-training map and a PCP5 map. The algorithm began by computing the average time-in-position map from the first 15 minutes of all the preinjection recordings. Step 2 used the average pre-injection time-in-position map to randomly select a subset of samples from the 30-min sessions in which the moderate dose of PCP was injected. The goal of this step was to down-sample the PCP5 sessions and create the best approximations from the first 30 minutes of the PCP sessions to the average 15-min pre-injection time map. These approximations were relatively poor. Thus the third step computed the average time map from the set of PCP5 time maps that were created in step 2. The final step used the average down-sampled PCP5 map to randomly select samples from all the 30-min data time series. As a result, we selected data from each recording using the same algorithm so that the time-in-position maps were similar in all the recordings we would compare. These time-in– position maps accounted for a pseudo-random, unbiased ~15-min sample from throughout the 30-min recording. Examples of the average time-in-position maps for each type of session (preinjection, PCP0, PCP3, PCP5) are shown in Fig. 5A. The measures of place cell quality were computed from the standardized recordings.

### Electrophysiological data analysis - Overdispersion

The overdispersion in place cell discharge is computed as the variance of the standardized firing rates of the cell for passes through the firing field, as previously described (Fenton et al., 2010). The standardized firing rate (*z*) was computed for each 5-s interval as:

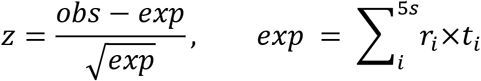

where *obs* is the number of observed action potentials and *exp* is the number of action potentials that were expected according to the assumption of Poisson firing. The expectation is the sum of the products of the time spent (*t*_*i*_) in a location during time interval *i* and the time-averaged rate at that location (*r*_*i*_) computed for all locations that were visited during the 5 seconds. Note that if the rat does not visit places where the cell fired during the entire recording, then *exp* = 0 and *z* is undefined. To select intervals when the rat sampled a cell’s firing field well, only those 5-s intervals were selected for the computation where *exp* was greater than or equal to five times the average rate for the cell.

### Electrophysiological data analysis - Functional coupling of cell pairs

The functional coupling of spike trains from pairs of cells was estimated using Kendall’s correlation (Press et al., 1993), which was computed as previously described (Olypher et al., 2006). The correlation was computed between the time series of spike counts from the two cells. The time series was generated by counting the number of spikes the cell fired during each 250-ms interval.

### Electrophysiological data analysis - Correlation of ensemble activity across time

The action potential discharge within an ensemble of *n* cells during a particular time interval can be characterized by an activity vector of dimension *n*, which is just a list of spike counts that describes the number of spikes emitted by each cell during the interval (see example Fig. 10B). The similarity of the ensemble’s activity between two time intervals was calculated as Pearson’s correlation of the two vectors. We computed the set of ensemble activity correlations between all pairs of 1-min time intervals during a recording. The set of correlations were organized and displayed as a correlation matrix (e.g. Fig. 10B). Correlations during a range of time intervals were quantified by computing the average correlation in the corresponding region of interest of the matrix.

### Electrophysiological data analysis - Estimating the ensemble activity due to changes in behavior

The ensemble activity correlations were dramatically changed by PCP injections (Fig. 10F). The change could arise because PCP disturbed the temporal discharge relationships of the cells as predicted by the discoordination hypothesis, or alternatively, because PCP disturbed the rat’s behavior, such that the sampling of the environment was distinct in the pre- and post-injection intervals. We evaluated this behavioral possibility by simulating ensemble spike trains on the basis of the pre-injection firing rate maps and the rat’s current behavior, as previously described (Fenton et al., 2010). The resulting Poisson spike trains were evaluated exactly like the raw data and results of analyzing the simulation and the real data were compared (Fig. 11).

### Electrophysiological data analysis - Decoding position from ensemble discharge

Estimates of the rat’s location (*x*) were computed from the single unit ensemble activity vector (*n*), using a published Bayesian algorithm (Zhang et al., 1998), where the posterior probability of the current location is defined as 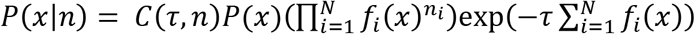, where *C*(*τ*,*n*) is a normalization factor so that ∑_*x*_ *P*(*x*|*n*) = 1, *f*_*i*_(*x*) are firing rate maps for cells *i*..*N* obtained by binning the 2-D space into 32×32 2.5 cm bins, *P*(*x*) is the time-in-location dwell distribution, *τ* is the length of the time window (300 ms), *n*_*i*_ is the number of spikes fired by the i-th cell in a given time window and *x* is the *(x,y*) position of the rat. Intervals with no spikes were excluded from analysis. Centered posterior probability plots as in Fig. 6, are computed by shifting the posterior probability such that the rat’s observed location is at the center of the plot. To obtain the decoded location estimate, we selected the 10% of locations with the largest posterior probability and computed the probability-weighted average of their x and y coordinates. The decoding error was computed as the Euclidean distance between the observed and estimated locations. Confidence in the decoding was estimated as the “Posterior confidence area,” which is the area where the posterior is reliably elevated. The threshold was preset for asserting the probability of locating the rat was elevated. The threshold was obtained independently in each pre-injection recording as the locations with posterior probability greater than the mean+std probability.

### Experimental Design and Statistical Analysis

Test statistics, the associated degrees of freedom (d.f.), and p values are reported. Analytical values are given to three significant digits. Statistical calculations were performed using JMP 12, Matlab versions 8.5 and higher, and custom software written in C. Power analyses used GPower 3.0.6 (Franz Faul, Edgar Erdfelder, Albert-Georg Lang, and Axel Buchner).

Statistical design for analysis of the active place avoidance behavioral experiments can be found in the Results that describe Figure 2C-G. Briefly, a within-subjects design was used to assess the effect of systemic PCP on behavior and hippocampus cell firing in the awake freely-behaving rat. Baseline measurements were made and then PCP and control solutions were administered in pseudo-random order with at least a 2 d wash out period. This period was judged to be adequate because PCP-induced alterations could not be detected in neural activity or behavior beyond 1 h after treatment. Changes in behavior after the different treatments were compared.

Statistical design for analysis of the LFP electrophysiological recordings during the behavioral tests can be found in the Results that describe Figure 2H.

Statistical design for analysis of the LFP electrophysiological recordings during foraging for scattered food that were examined for the changes in LFP power, phase-amplitude comodulation, phase and amplitude features of gamma oscillations can be found in the Results that describe Figure 3.

**Figure 3.**
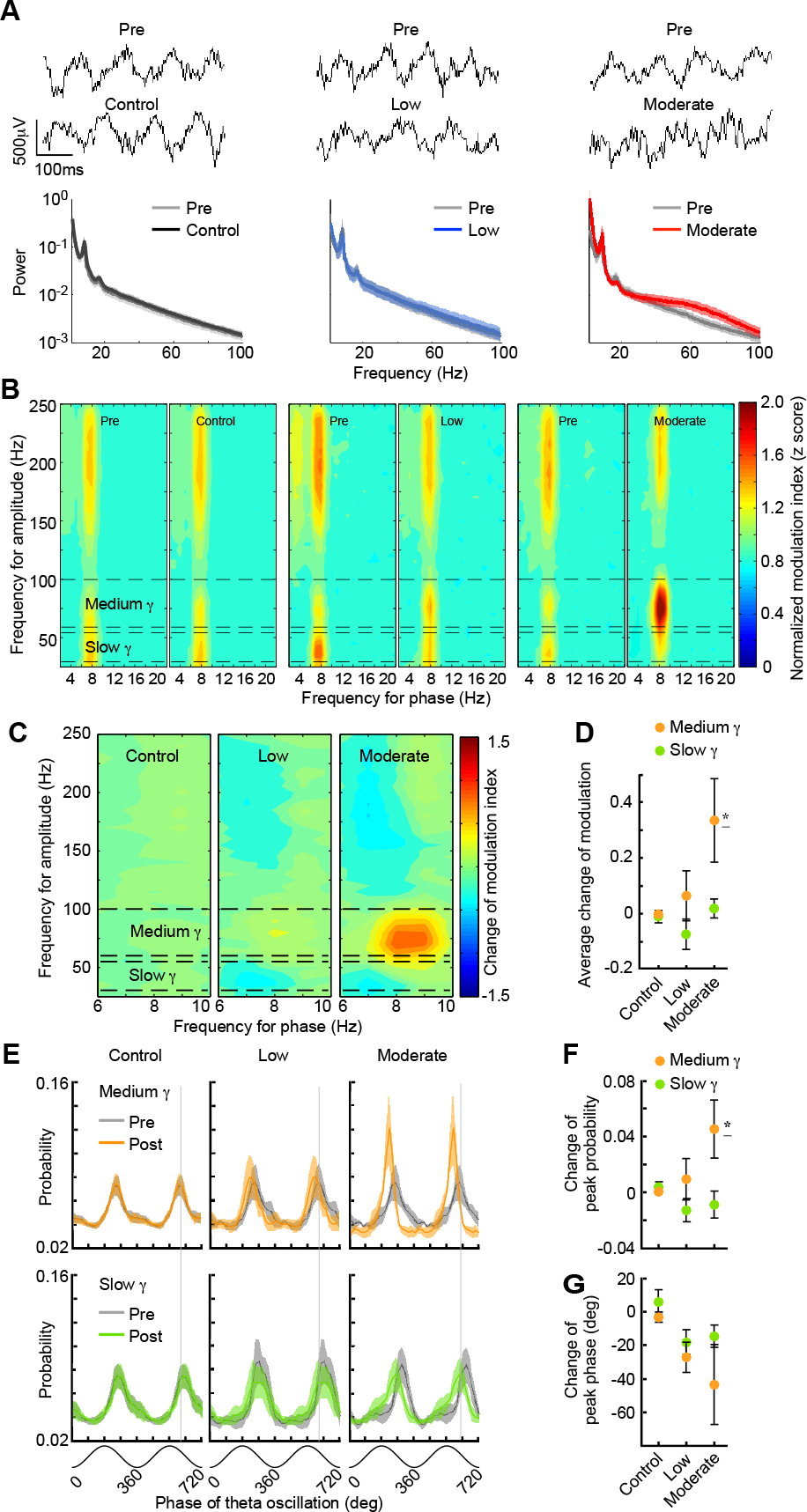
PCP discoordinates CA1 local field potentials (LFPs). (A) Example LFPs at *stratum pyramidale* and the average power spectra from the 30-min before and after systemic PCP administration. Power is relative to the maximum of all six spectra. (B) Pre- and post-treatment averaged cross-frequency coupling measured by comodulograms and (C) the average post minus pre-treatment difference comodulograms. The changes are quantified in (D) by average PCP-induced changes of cross-frequency coupling between theta and slow gamma (30-55 Hz), and theta and medium gamma (60-100 Hz) oscillations. (E) Average theta-phase modulation profiles of slow and medium gamma oscillations. Average effect of PCP administration on (F) the peak probability and (G) the phase of theta-modulated slow and medium gamma oscillations. Error bars and shading: ± S.E.M.; Pre: Pre-injection; Post: Post-injection; Control, low, moderate: 0, 3, 5mg/kg PCP doses. 12, 4, 5 recordings in control, low, and moderate, respectively. <u>* indicates</u> p < 0.05 relative to all groups.

Statistical design for analysis of the CA1 single unit electrophysiological recordings during foraging for scattered food focused on the firing rate, spatial coherence, information content, map similarity and probability of firing during passes through the firing field. The design can be found in the Results that describe Figure 4. The design for analysis of these data after standardizing the position sampling can be found in the Results that describe Figure 5. Statistical design of the analysis of these data to assess accuracy and uncertainty of decoding the rat’s position can be found in the Results that describe Figure 6.

**Figure 4.**
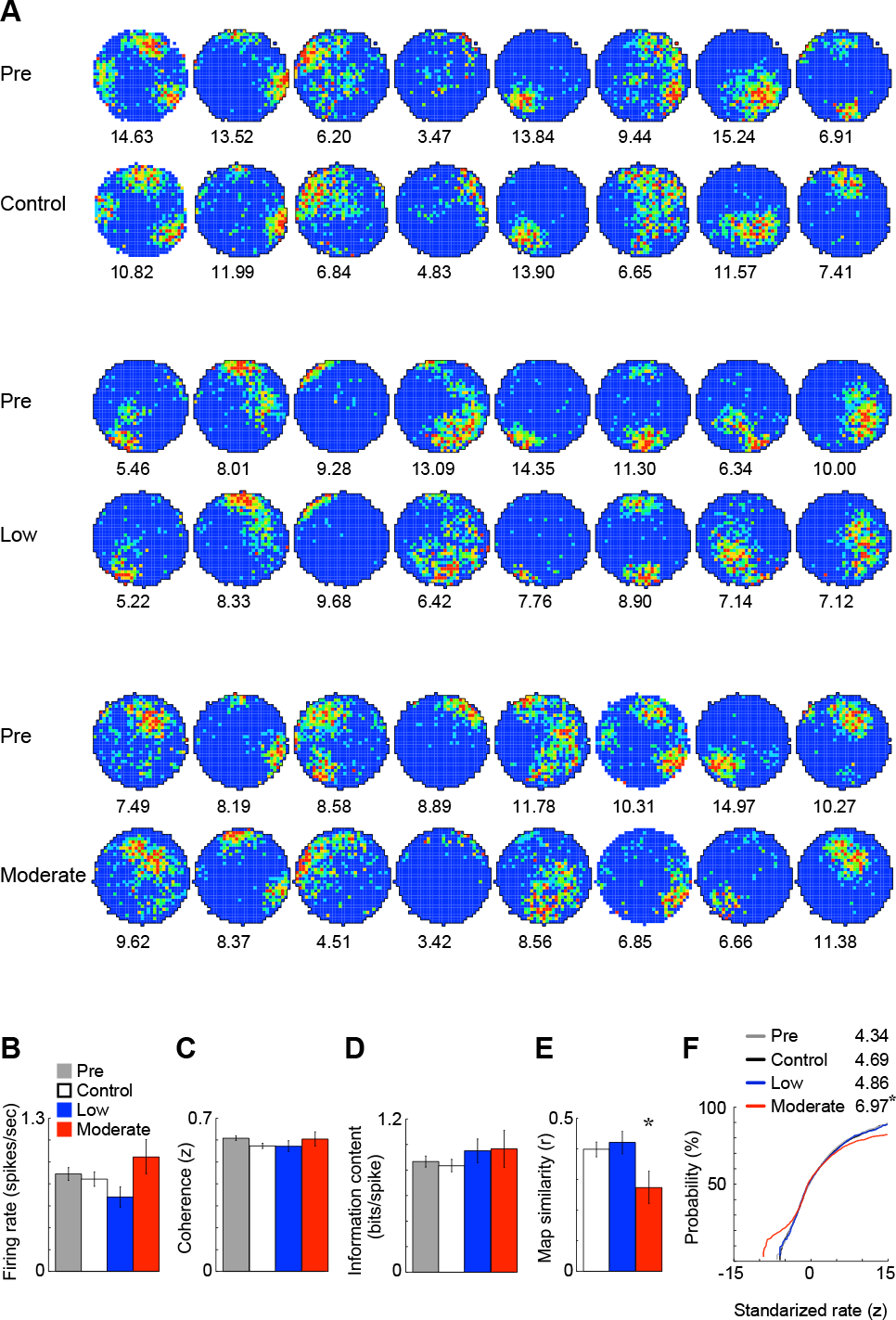
Place cell discharge properties are preserved after systemic PCP administration. (A) Blue-to-red color-coded firing rate maps from 8 example place cells recorded before and after control, low, and moderate doses of PCP. The median firing rate in the maximum-rate (red) category is given below each map. Pre- and post-injection measures of the quality of place cell discharge were unchanged by the PCP injections: (B) Firing rate; (C) Spatial coherence; (D) Spatial information content. (E) The moderate dose of PCP affected the pre-post session similarity of the firing rate maps. (F) The moderate dose of PCP increased the overdispersion in place cell discharge. The plots are the cumulative probability distribution of standardized firing rates during 5-sec episodes when the rat traversed a place cell’s firing field. Overdispersion measured as the variance of the distribution is given. Error bars: ± S.E.M.; Comparisons were made by one-way ANOVA; Pre: Pre-injection; Control, low, moderate: 0, 3, 5mg/kg PCP doses. 64 place cells are recorded from 12 recordings in control, 26 from 4 recordings in low, and 17 from 5 recordings in moderate PCP dose. * indicates p < 0.05 relative to all groups.

**Figure 5.**
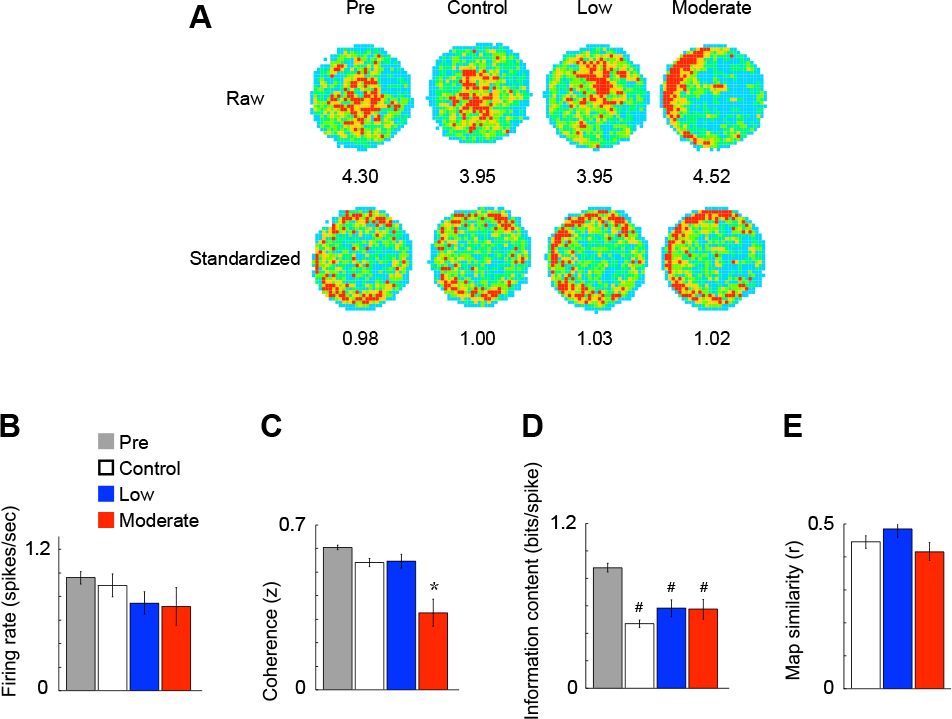
Standardization of position sampling. (A) The location-specific data for evaluating place cell firing were standardized to eliminate differences in spatial sampling caused by systemically-administered PCP-induced movement stereotypy. Blue-to-red color-coded averaged time-in-location maps before (upper row) and after (bottom row) standardization. After the moderate dose of PCP, the rat spent more time at the periphery of the arena due to the PCP-induced stereotypy. After standardized position sampling (bottom row), the post-injection time-in-location maps for each PCP dose were more similar than before standardization (upper row). Pre- and post-injection measures of the quality of place cell discharge after standardization: (B) Firing rate (F_3, 338_ = 1.57, p = 0.196); (C) Spatial coherence (F_3, 338_ = 23.98, p = 4.29×10^−14^); (D) Spatial information content (F_3, 338_ = 24.67, p = 1.85×10^−14^); (E) Similarity of the firing rate maps (F_2, 132_ = 1.99, p = 0.141). Error bars: ± S.E.M.; Comparisons were made by one-way ANOVA; Pre: Pre-injection; Control, low, moderate: 0, 3, 5mg/kg PCP doses. * indicates p < 0.05 relative to all groups, # indicates p < 0.05 relative to pre-inject group.

**Figure 6.**
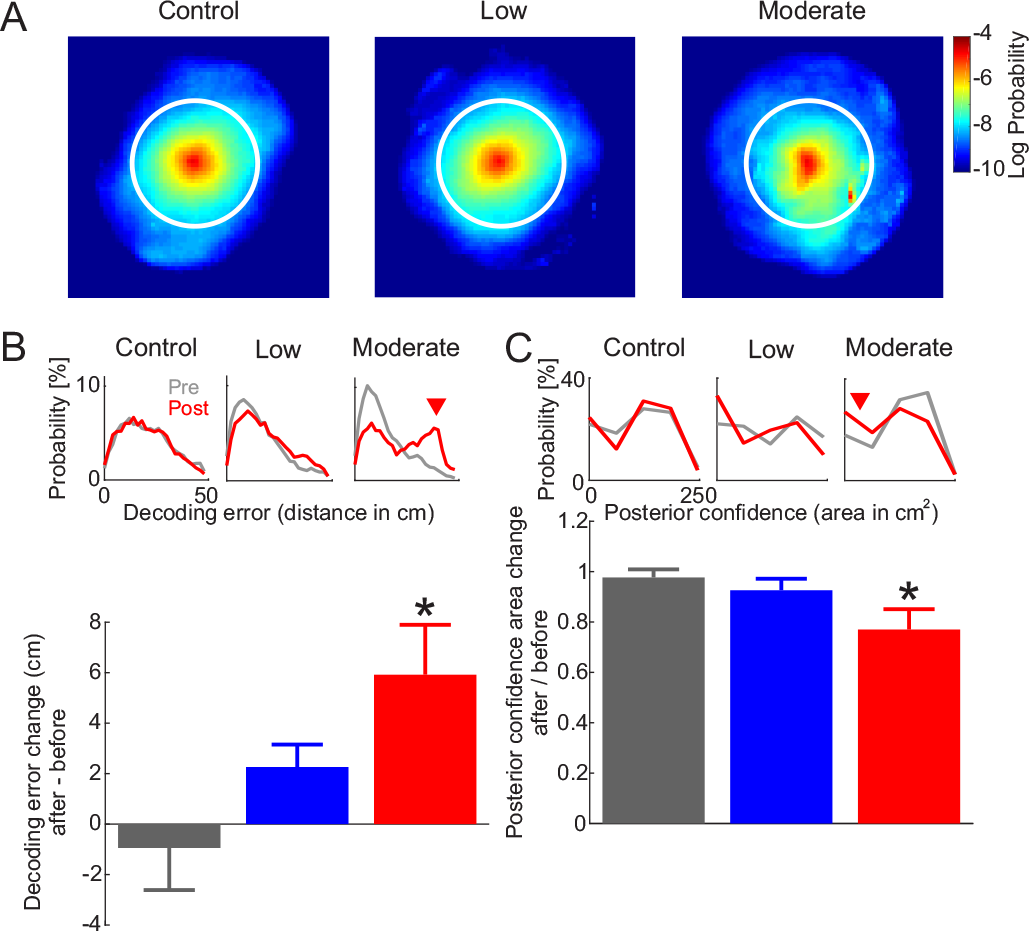
The moderate systemic dose of PCP decreases the accuracy and increases the uncertainty of the rat’s position decoded by Bayesian comparisons of post-injection ensemble discharge during 300 ms epochs to the probability of observing pre-injection spatial ensemble discharge. (A) Average decoded posterior probabilities centered at the rat’s actual location for 0mg/kg (Control), 3mg/kg (Low) and 5mg/kg (Moderate) PCP doses. The white circle indicates the size of the circular arena. (B) Top: Example distributions of the decoding error for pre- (gray) and post-injection (red) recordings showing both accurate decoding and increased decoding errors after the moderate PCP dose (red triangle). Bottom: Average decoding error for Control (gray), Low (blue) and Moderate (red) PCP doses. (C) Top: Examples distributions of the decoding uncertainty, estimated by the elevated posterior area for pre- (gray) and post-injection (red) recordings showing increased uncertainty after the moderate PCP dose (red triangle). Bottom: Comparison of the average uncertainty of the decoding between pre- and post-injection for Control (gray), Low (blue) and Moderate (red) PCP doses. Error bars: ± S.E.M.; Comparisons were made by one-way ANOVA * indicates p < 0.05 relative to Control.

The statistical design for the combined analysis of single unit recordings and LFPs using their phase-frequency discharge preference can be during foraging for scattered food can found in the Results describing Figure 7. The statistical design for analysis of these data to assess the discharge coordination of place cells can be found in the Results describing Figure 8 and the statistical design for assessing the change of discharge coordination by evaluating pairs of simultaneously recorded place cells can be found in the Results describing Figure 9A. Similarly, the statistical design for assessing the change of discharge coordination by evaluating pairs of simultaneously recorded single units during restriction to a small environment can be found in the Results describing Figure 9B.

**Figure 7.**
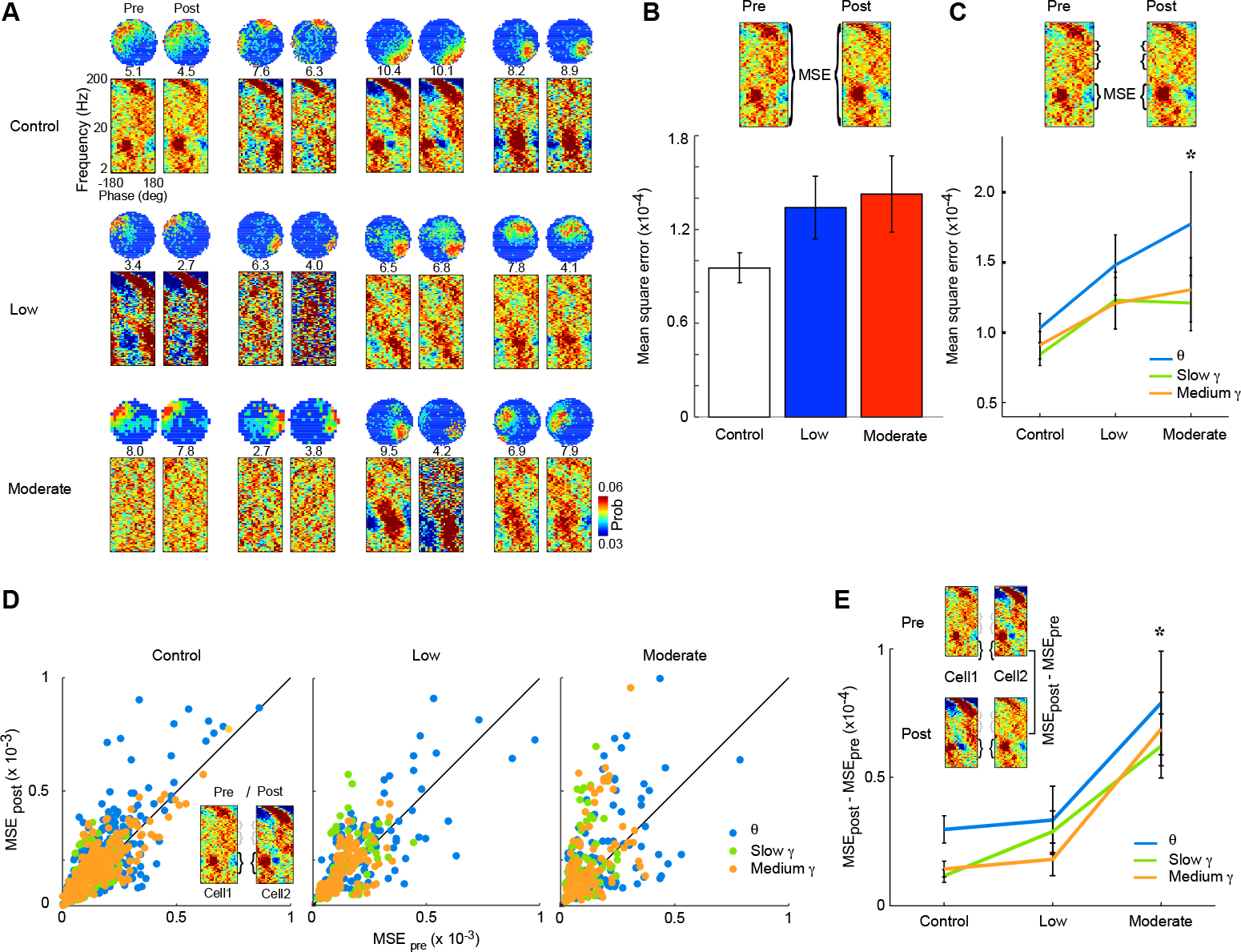
PCP changes discharge phase-frequency preference of principal cells. (A) Phase– frequency-discharge probability plots for example place cells recorded before and after systemic PCP injection. Probability in the plot is indicated by color and the firing rate map of the cell is given above. Stability of (B) wideband and (C) band-specific phase-frequency-discharge probability after PCP treatment. (D) Scatter plots and (E) summary plots of the cell-pair stability in the relationship between the individual phase-frequency-discharge probability maps before and after PCP. Error bars: ± S.E.M.; Comparisons were made by ANOVA; Pre: Pre-injection; Post: Post-injection; Control, low, moderate: 0, 3, 5mg/kg PCP doses. 64 place cells are recorded from 12 recordings in control, 26 from 4 recordings in low, and 17 from 5 recording in moderate doses of PCP. * indicates p < 0.05 relative to control.

**Figure 8.**
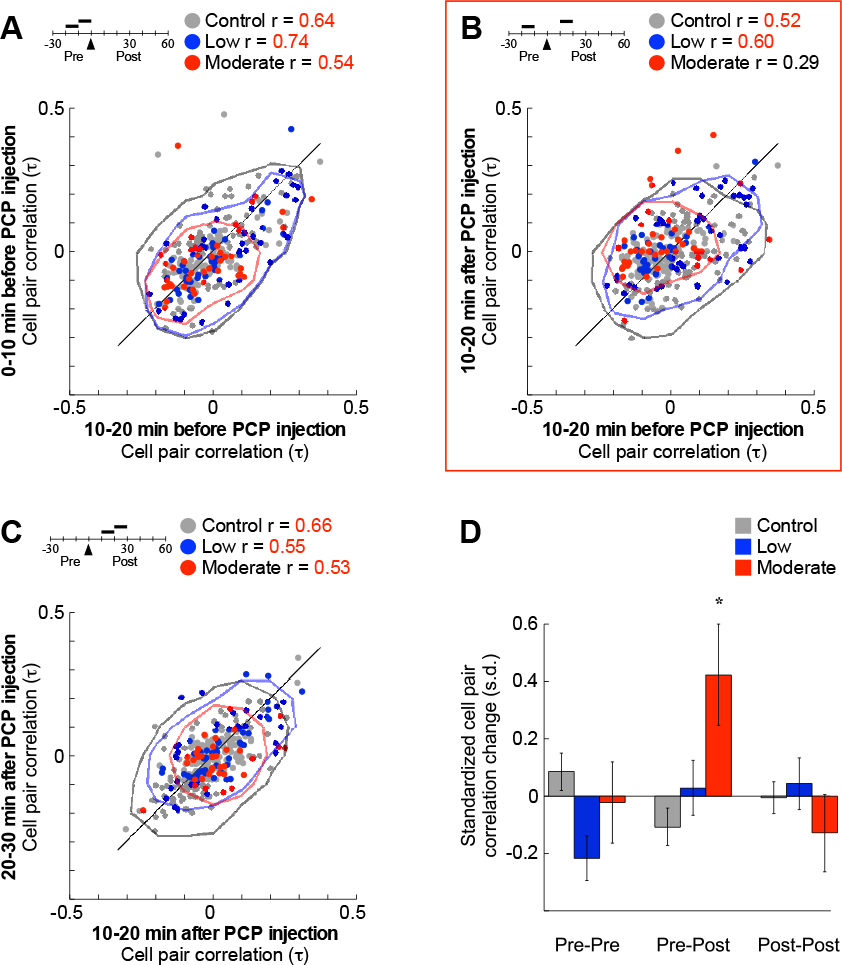
Systemic PCP induced instability of sub-second time scale discharge coordination of place cells. Cell-pair coordination was estimated by computing Kendall’s correlation (τ), for all pairs of simultaneously recorded place cells. Spike counts were determined each 250 ms and the correlation was computed during 10-min epochs of the pre- and post-injection recordings. Stability of the coordination is estimated by PCo (Lee et al., 2014), the Pearson correlation (r) comparing cell-pair coordination in two epochs for (A) two pre-injection epochs; (B) pre- and post-injection epochs and (C) two post-injection epochs. The iso-density contours in A-C highlight the relative changes in cell-pair coordination. (D). The change of coupling between the discharge of cell pairs in A-C was normalized as: (τ_late_ - τ_early_)/standard deviation [early]. A total of 206 place cell pairs were examined after the control dose, 75 pairs after the low dose, and 47 pairs after the moderate dose. Error bars: ± S.E.M.; Comparisons were made by ANOVA; Significant (p < 0.05) Pearson correlations are indicated by red. * indicates significantly different from all others.

**Figure 9.**
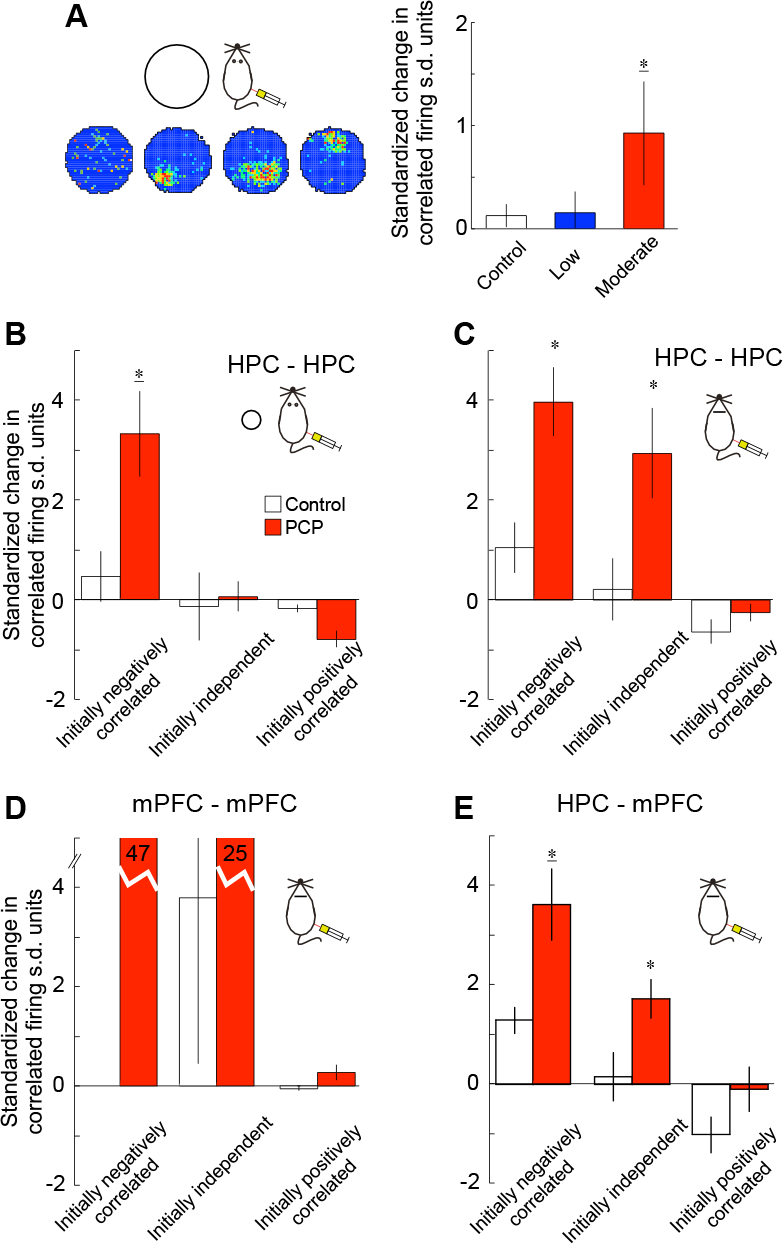
Systemic PCP alters neural coordination independently of hyerlocomotion. (A) The change in Kendall’s correlation (τ) from before to after PCP treatment was computed for all pairs of simultaneously recorded complex-spike cells, not just place cells (4 examples shown) as rats foraged in the cylinder. For each cell pair, spike counts were determined each 250 ms and we computed the average correlation during the six 5-min episodes from the 30-min pre- and postinjection recordings. Pre: Pre-injection; Control, low, moderate: 0, 3, 5mg/kg PCP doses. * p < 0.05 relative to control. (B and C) PCP changes the coupling of the cell-pairs that are negatively correlated before injection, but not the cell-pairs that were positively correlated before injection. The potential influence of location-specific place cell firing and (6 mg/kg i.p.) PCP-induced hyperactivity was minimized by recording rats (B) that were restricted to a 24-cm diameter circular enclosure with 20-cm high walls and by recording rats (C-E) under urethane anesthesia. The average and standard deviation (s.d.) of these pre-injection and post-injection episodes were computed for each cell pair. Cell pairs with significant negative or positive pre-injection correlations were categorized as “initially negatively correlated” or “initially positively correlated” and those that were not significant correlated were categorized as “initially independent.” Next, for each cell pair we computed the standardized change of the correlation after each PCP dose. The standardization was done according to: (post - pre)/s.d.(pre), where pre and post indicate the average pre-injection and post-injection correlations, respectively. This standardization was done to avoid the bias that can arise from estimating proportional changes in small numbers, such as correlations near zero. The moderate dose of PCP selectively increased the coupling of only those cell pairs that were negatively or independently coupled prior to PCP administration, without affecting the coupling of initially positively correlated cell pairs. Error bars: ± S.E.M.; Comparisons were made by ANOVA; * indicates p < 0.05 relative to the control; <u>* indicates</u> p < 0.05 relative to all other groups.

The statistical design for the electrophysiological single unit recordings under anesthesia to assess the change of discharge coordination by evaluating pairs of simultaneously recorded single units can be found in the Results describing Figure 9C-E.

Statistical design to analyse the change of discharge coordination of place cells in the single unit recordings from CA1 during foraging for scattered food can be found in the Results describing Figure 10C-G. The statistical design for analysis of the impact of spatial sampling on these analyses can be found in the Results describing Figure 11.

**Figure 10.**
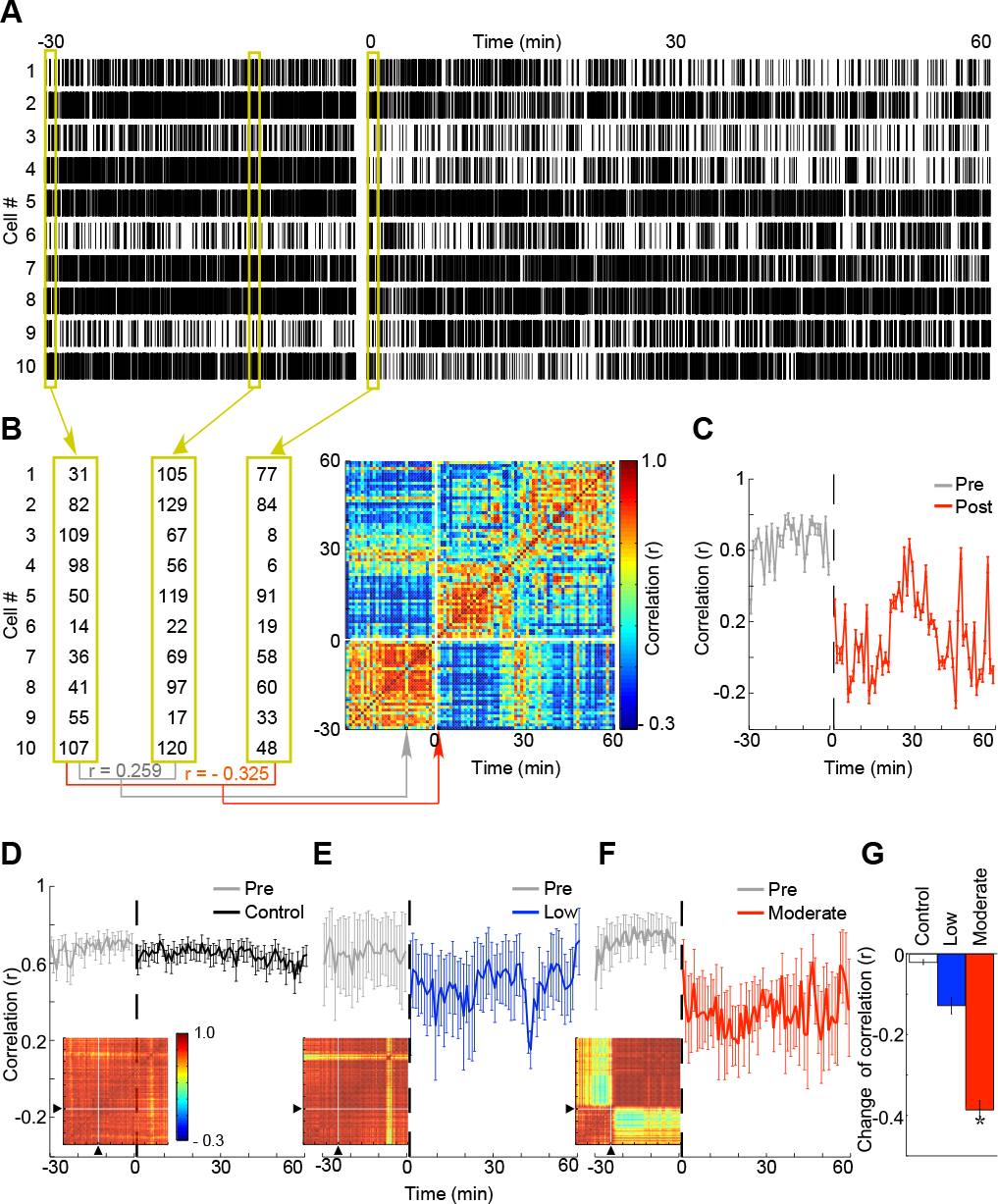
PCP induced instability of multi-second time scale discharge coordination of place cells. (A) Raster plots representing a 10-cell ensemble recording in which 5 mg/kg PCP was injected after 30 min, at time = 0. Ensemble activity vectors were computed each minute by summing the number of action potentials each cell discharged during the minute. Three of these minute-long activity vectors are illustrated in yellow rectangles. (B) Each activity vector was compared with every other activity vector using Pearson’s correlation coefficient (r) and the results are organized in the color-coded correlation matrix. (C) The correlation was quantified by computing the average correlation between the current activity vector and all the pre-injection vectors. The average ± S.E.M. of such correlations is shown for the different PCP doses (D) control (n = 12 ensembles), (E) low (n = 4 ensembles) and (F) moderate (n = 5 ensembles). (G) The average post- versus pre-injection change in the correlation quantifies that the moderate dose of PCP changed the ensemble activity discharge pattern to a new pattern that did not resemblethe hippocampal ensemble patterns before the * indicates p < 0.05 relative to all groups.

**Figure 11.**
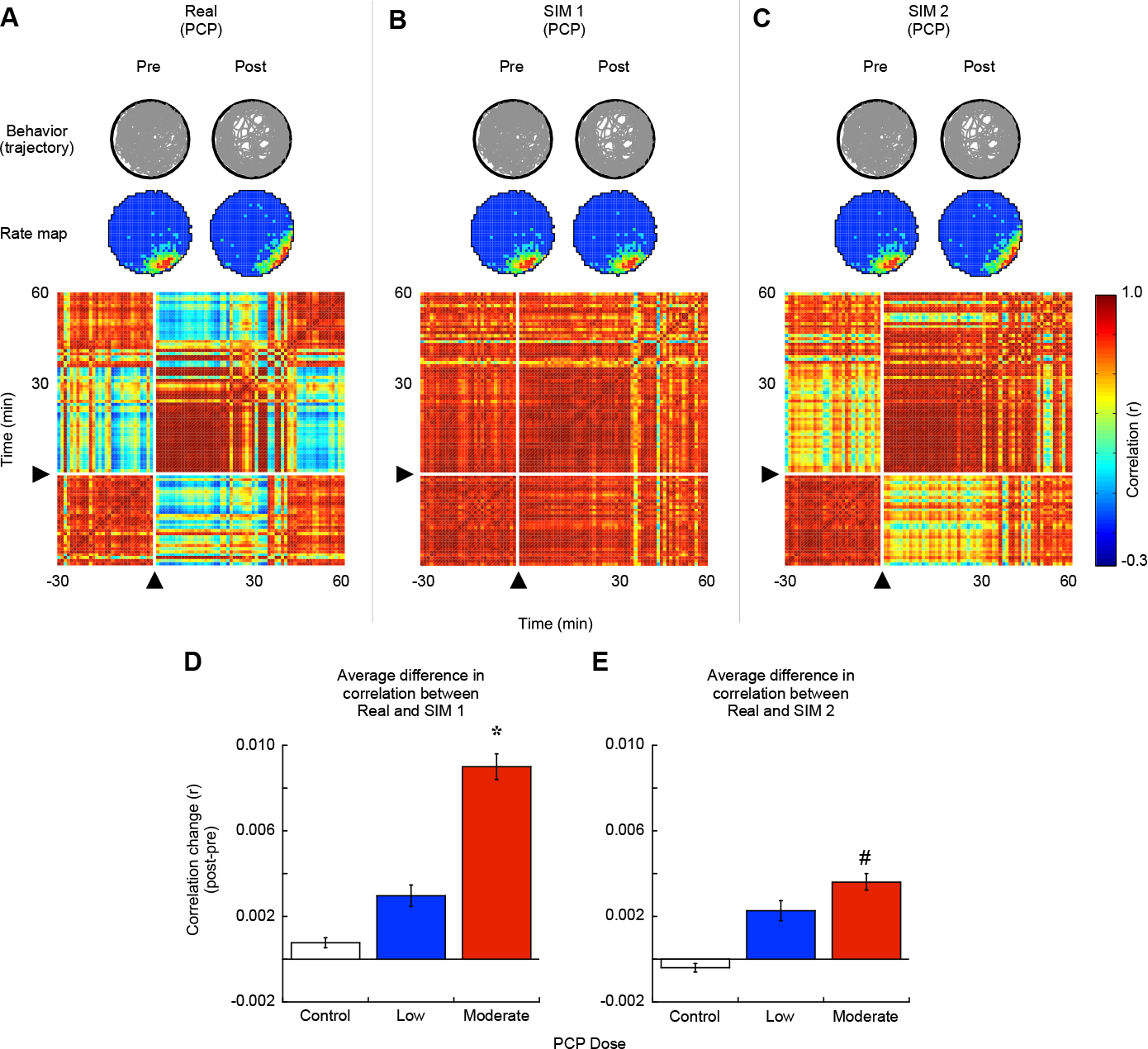
The PCP-induced neural discoordination is not explained by PCP-induced changes in motor behavior and subtle changes in firing rate maps. (A) Real data. Rat’s trajectory during pre- and post-injection sessions and example firing rate maps in pre- and post-injection sessions. The correlation matrix that compares all pairs of ensemble activity vectors for an example recording is shown. In this “Real” matrix, the moderate dose of PCP, was injected at time = 0. The change of ensemble activity pattern is manifested by decrease in correlation coefficients comparing pre- and post-injection activity. The source of this change in activity was tested using two simulations. (B) Simulation 1 (SIM 1). Rat’s trajectory and example firing rate map used in the simulation. The spike trains were simulated using an inhomogeneous Poisson model (e.g. Fenton et al. 2010) based on location-specific firing rate maps that were recorded before PCP injection and the actual spatial trajectory that the rat executed before and after PCP injection. The Poisson spike trains represent the location-specific discharge expected from the rat’s particular behavior each minute, assuming that the location-specific firing rate map did not change after the PCP injection. The correlation matrix created from simulated data. The change in ensemble activity, which is visible in real data, is not detected in the SIM 1 model, showing that change in the ensemble activity pattern can’t be accounted for by change in rat’s motor behavior. (C) Simulation 2 (SIM 2). Rat’s trajectory and example firing rate maps used in the simulation. The spike trains were simulated based on the location-specific firing rate maps that were recorded before and after PCP injection and the actual spatial trajectory that the rat executed before and after PCP injection. The correlation matrix created from simulated data. The change in ensemble activity after PCP injection in the SIM2 model is smaller than the change in activity observed in real data, showing that the change in ensemble activity pattern can’t be fully accounted for by the change in the rat’s motor behavior and changes in firing rate maps. (D) The average difference (±S.E.M.) in correlations between “Real” and “SIM 1” correlation matrices following control, low or moderate dose of PCP. The data compare the corresponding areas of the matrices that represent the post-injection activity vector correlations with the pre-injection activity vectors. The effect of PCP dose was significant (F_2,18_ = 13.17; p = 2.99×10^−4^, ANOVA) because moderate PCP differed from both control and low dose of PCP (post-hoc: control = low > moderate, p’s < 0.004). (E) The average difference (±S.E.M.) in correlations between “Real” and “SIM 2” correlation matrices following control, low or moderate dose of PCP. The data compare the corresponding areas of the matrices that represent the post-injection activity vector correlations with the pre-injection activity vectors. The effect of PCP dose was significant (F_2,18_ = 5.14; p = 0.171, ANOVA). Pre: Pre-injection; Post: Post-injection; Control, low, moderate: 0, 3, 5mg/kg PCP doses. * indicates p < 0.05 relative to all groups, # indicates p < 0.05 relative to control group.

Because neural activity parameters can strongly vary amongst cells and between cell types, the comparisons of treatment effects on neural activity were computed relative to the baseline values of the particular cell. Assessing the effect of PCP on neural activity in the anesthetized rat was based on a between-subjects design. Each subject received either a PCP or control solution. Differences between treatments, categories of cell pair, or time intervals were compared by Student’s t test, one-way and multiple-way ANOVAs, as appropriate and described in Results. Analysis of within-subjects design data used paired-t test and repeated measures ANOVA. When all pair-wise post-hoc comparisons were examined, Newman-Keuls post-hoc comparisons were performed. Dunnett’s test was used for post-hoc comparisons against the control value. Statistical significance was accepted when p < 0.05. All statistical comparisons of correlation values such as spatial similarity and coherence were performed on Fisher’s z transform of r:

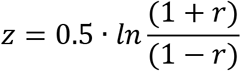

## RESULTS

### Cognitive behavioral impairment under PCP is indexed by excessive medium gamma oscillations

Using the active place avoidance paradigm (Fig. 2A), we began by confirming that PCP impairs performance of a familiar spatial navigation task that requires cognitive control (Kelemen and Fenton, 2010, 2016). Animals on a continuously rotating disk-shaped arena must use distal spatial cues to identify the location of a shock zone that is stable in the room. The rotation makes spatial information from the rotating disk irrelevant for avoiding the shock zone. This two-frame (‘Room+Arena-’) variant of the task challenges the subject to use relevant spatial information from the room spatial frame and ignore the irrelevant information from the rotating arena spatial frame (Fenton et al., 1998; Fenton and Bures, 2003; Wesierska et al., 2005), which we have used to operationally define cognitive control in prior work that directly measured that place information about location in the room and in the arena are alternatively being represented in place cell spike trains during active place avoidance (Kelemen and Fenton, 2010). A separate one-frame (‘Room+’) task variant was also tested. It requires avoidance of the same place in identical conditions except that the arena surface is covered by shallow water to reduce the salience of arena cues that are irrelevant for avoiding shock (Fig. 2B). The shallow water reduces the demand for cognitive control to ignore the irrelevant arena information while using the relevant room information to avoid the constant current foot shock (Wesierska et al., 2005; Lee et al., 2012).

Well-trained rats (n = 10) received a systemic injection of control (0 mg/kg), low (3 mg/kg), and moderate (5 mg/kg) doses of PCP 30 min before testing (Fig. 2C inset). PCP caused the well-known dose-dependent increase in locomotion (Fig. 2C left panel; F_2,27_ = 9.94, p = 5.82×10^−4^, ANOVA, post hoc tests: control = low < moderate). However, only the moderate dose impaired the familiar two-frame active place avoidance measured by the number of entrances to the shock zone (Fig. 2D left panel; F_2,27_ = 6.11, p = 6.47×10^−3^, ANOVA, post-hoc tests: control = low < moderate). The impairment could be due to PCP reducing the sensitivity to shock, but this possibility is inconsistent with the observation that when the rats entered the shock zone, they escaped even faster at all three PCP doses, measured by the change in the average number of shocks they received during each entrance (Fig. 2E left panel; F_2,26_ = 0.03, p = 0.970, ANOVA; one trial had no entrances). These data suggest that PCP impairs expression of a conditioned place avoidance memory as well as induces hyperlocomotion.

Could the cognitive impairment be due a PCP-induced disregard for the shock or inability to perceive the relevant stimuli, or the concomitant hyperlocomotion, as we have shown in a different rat model (O’Reilly et al., 2016)? We observed that the moderate dose of PCP also induced hyperlocomotion in the Room+ task variant (Fig. 2C right panel; n = 4 rats; t_3_ = 4.29, p = 0.023, paired t-test), compared to control saline injections. However PCP did not impair the same conditioned place avoidance in this task variant with low cognitive control demand (Fig. 2D right panel; t_3_= 0.67, p = 0.551, paired t-test). Again, PCP did not increase the number of shocks that were received once the rat entered the shock zone (Fig. 2E right panel; t_3_ = 1.01, p = 0.387, paired t-test).

In a further effort to disentangle the hyperlocomotion effects of PCP from its potentially confounding effect on cognitive performance (Moghaddam and Adams, 1998), we tested whether intrahippocampal infusion of PCP is sufficient to reproduce the systemically-induced impairment of familiar active place avoidance (Fig. 2F inset). Compared to bilateral intrahippocampal saline infusions, PCP infusions of 6,8 or 10 μg/μl/side, did not induce hyperlocomotion (Fig. 2F; t_22_ = 1.89, p = 0.072, paired t-test). Nonetheless, PCP impaired place avoidance measured as an increased number of entrances (Fig. 2G; t_22_ = 2.47, p = 0.022, paired t-test). In the subset of eight rats from which we also recorded the hippocampal local field potential (LFP), the magnitude of the PCP-induced performance deficit was significantly related to the magnitude of the change in CA1 medium gamma (60-100 Hz) oscillation amplitude (Pearson’s correlation coefficient r = 0.94, df = 6, p = 1.63×10^−3^), but not to the change in theta (Pearson’s correlation coefficient r = 0.32, df = 6, p = 0.484) or slow gamma (Pearson’s correlation coefficient r = 0.42, df = 6, p = 0.348) oscillation amplitudes (Fig. 2H). These findings demonstrate that PCP in dorsal hippocampus is sufficient to impair learned place responding in a task with significant cognitive control demand although potentially confounding hyperlocomotion was absent. The findings also demonstrate that PCP-induced increases in hippocampus medium gamma oscillations predict the magnitude of the cognitive behavioral impairment, a novel finding that is predicted by the hypothesis that dominance of CA1 medium gamma over slow gamma, biases hippocampus output to process information currently available to the senses at the expense of what has been learned (Colgin et al., 2009; Fries, 2009; Keeley et al., 2016). All subsequent experiments examined the effects of systemic PCP administration.

### Aberrant local field potential oscillations under PCP

Next we analyzed the effect of systemically administered PCP on the LFP at CA1 *stratum pyramidale*. These recordings were concurrent with the CA1 place cell recordings we analyze next. CA1 LFPs were reliably altered for 30 minutes after administering the moderate dose of PCP so we compared LFPs during the 30 minutes before and after drug administration (Fig. 3A). Only after the moderate PCP dose, CA1 gamma became exaggerated (F_2,18_ = 4.33, p = 0.029, ANOVA; Fig. 3A). Delta oscillations (1-4 Hz) also increased (F_2,18_ = 10.64, p = 9.68×10^−4^, ANOVA) but neither theta (4-12 Hz; F_2,18_ = 2.03, p = 0.160, ANOVA) nor beta (12-30 Hz; F_2,18_ = 1.21, p = 0.348, ANOVA) oscillations changed (Fig. 3A).

We then investigated the coordination of theta and gamma oscillations, as has been reported in human (Canolty et al., 2006), rat (Tort et al., 2009; Dvorak and Fenton, 2014) and mouse (Radwan et al., 2016) subjects. Prior to PCP, theta-gamma phase-amplitude coupling was stronger for slow gamma (30-55 Hz) than for medium gamma (60-100 Hz) oscillations. After the moderate PCP dose, the theta-phase modulation of medium gamma oscillations increased significantly without changing the modulation of slow gamma oscillations (Fig. 3B-D). Both the slow and the medium gamma oscillations occurred at the descending phase of the theta oscillations with medium leading slow (Fig. 3E). The moderate dose of PCP changed the medium gamma modulation in two ways without affecting the modulation of slow gamma (treatment x phase interaction: F_2,36_ = 4.47; p = 0.019, ANOVA; all post-hoc comparisons to medium gamma after moderate PCP p’s < 0.01). In particular, the peak of the phase distribution increased after PCP administration (post-hoc p < 0.05; Fig. 3F) and increasing doses of PCP shifted the preference of the medium gamma oscillations to earlier phases of theta (F_2,18_ = 3.6; p = 0.048, ANOVA; Fig. 3G). Thus, after the moderate dose of PCP, the preferred theta phase of medium gamma oscillations became more phase specific with a preference for earlier phases, when pyramidal cell firing is normally less likely (Fox et al., 1986). PCP did not change the relationship between the theta-phase coupling of 100-250 Hz high frequency oscillation amplitudes called the fast gamma or epsilon band (Belluscio et al., 2012), a component of which is the discharge afterpotential, making this an indirect estimate of the action potential discharge rate within the CA1 network (Schomburg et al., 2012; Dvorak and Fenton, 2014). Accordingly, the lack of a PCP effect on theta-coupled 100-250 Hz oscillations predicts that PCP has no systematic effect on place cell discharge rates.

### Normal place field discharge properties of individual CA1 place cells under PCP

We then investigated the effect of PCP on the spatial discharge of place cells (n = 207; Fig. 4A) recorded from 4 rats before and after systemic administration of the control, low, and moderate doses of PCP. This provides a direct assay of how PCP affects a fundamentally cognitive variable, the neural representation of space. The average spatial discharge properties of place cells, specifically the firing rate (F_3,203_ = 1.60, p = 0.191, ANOVA; Fig. 4B), coherence (F_3,203_ = 1.79, p = 0.150, ANOVA; Fig. 4C), and information content (F_3,203_ = 0.67, p = 0.571, ANOVA; Fig. 4D) were not different after the injections. The similarity of firing rate maps before and after injection was lower after the moderate dose of PCP than it was after the other doses (F_2,96_ = 13.86, p = 5.15×1c^-6^, ANOVA; Fig. 4E), which was also reported for other NMDAR antagonists (Kentros et al., 1998; Ekstrom et al., 2001). However, this could be explained by the abnormal spatial sampling caused by PCP-induced hyperactivity and stereotypy (Fig. 2) because when the data were standardized for more homogeneous sampling, the PCP-induced difference in firing field similarity was no longer apparent (Fig. 5; Table 1). Standardizing the spatial sampling caused the spatial coherence of firing fields to decrease after the moderate dose of PCP (Fig. 5C) and a general decrease in the information content except for the pretraining session (Fig. 5D), demonstrating the impact of spatial sampling on firing field characteristics (Muller et al., 1987; Maurer et al., 2005). We also examined the pre-post injection differences of these discharge properties for individual cells. The absolute difference in each discharge property was higher after the moderate dose of PCP, indicating that this dose could modestly improve or degrade the spatial firing properties of individual place cells without causing a systematic change of the properties across the ensemble; PCP also did not alter the discharge properties of non-spatial presumptive pyramidal cells (n = 161) nor did it alter firing rates in the small sample of presumptive interneurons that were recorded (n = 11; Table 2). This overall preservation of place cell firing is consistent with reports that NMDAR antagonists do not disrupt place cell firing in familiar environments (Kentros et al., 1998; Ekstrom et al., 2001).

**Table 1.** Comparisons of CA1 place cell properties between raw and standardized data. After standardization, more cells meet the criteria to be classified as place cells. Data are represented as average ± S.E.M. Significant effects were confirmed by one-way ANOVA; * indicates p < 0.05 relative to all groups, # indicates p < 0.05 relative to control group.

| Original | Pre (n=100) | Control (n=64) | Low (n=26) | Moderate (n=17) |
| --- | --- | --- | --- | --- |
| Firing Rate (Spikes/sec) | $0.89 \pm 0.06$ | $0.84 \pm 0.07$ | $0.68 \pm 0.09$ | $1.05 \pm 0.16$ |
| Spatial Coherence (z) | $0.61 \pm 0.01$ | $0.57 \pm 0.01$ | $0.57 \pm 0.02$ | $0.61 \pm 0.03$ |
| Spatial Information Content (bits/AP) | $0.87 \pm 0.04$ | $0.83 \pm 0.05$ | $0.95 \pm 0.09$ | $0.97 \pm 0.14$ |
| Map Similarity (r) | | $0.40 \pm 0.02$ | $0.42 \pm 0.04$ | $0.27 \pm 0.05$ * |

Table 1. Comparisons of CA1 place cell properties between raw and standardized data.
| Standardized | Pre (n=171) | Control (n=82) | Low (n=39) | Moderate (n=50) |
| --- | --- | --- | --- | --- |
| Firing Rate (Spikes/sec) | $0.96 \pm 0.05$ | $0.89 \pm 0.10$ | $0.74 \pm 0.10$ | $0.72 \pm 0.16$ |
| Spatial Coherence (z) | $0.61 \pm 0.01$ | $0.54 \pm 0.02$ | $0.55 \pm 0.03$ | $0.33 \pm 0.06^*$ |
| Spatial Information Content (bits/spike) | $0.88 \pm 0.03$ | $0.47 \pm 0.03^{\#}$ | $0.58 \pm 0.06^{\#}$ | $0.58 \pm 0.07^{\#}$ |
| Map Similarity (r) | | $0.45 \pm 0.02$ | $0.49 \pm 0.03$ | $0.42 \pm 0.03$ |

**Table 2.** The CA1 non-spatial pyramidal cell and interneuron firing rate. No significant effects were confirmed by one-way ANOVA for both non-spatial pyramidal cells (F_3,157_ = 0.14, p = 0.936) as well as interneurons, although the presumed interneuron sample size is low (F_2,8_ = 0.93, p = 0.433). Data are represented as Average ± S.E.M.

| Non-spatial<br>pyramidal cells | Pre (n=79) | Control (n=48) | Low (n=15) | Moderate (n=19) |
| --- | --- | --- | --- | --- |
| Firing Rate<br>(Spikes/sec) | $0.73 \pm 0.06$ | $0.75 \pm 0.08$ | $0.64 \pm 0.12$ | $0.73 \pm 0.12$ |
| Interneurons | Pre (n=5) | Control (n=3) | Low (n=0) | Moderate (n=3) |
| Firing Rate<br>(Spikes/sec) | $12.43 \pm 3.33$ | $11.64 \pm 3.48$ | None | $6.48 \pm 1.26$ |

Place fields were generally preserved under PCP indicating spatial reliability, so we examined overdispersion, which estimates the temporal reliability of location-specific place cell discharge (Fenton and Muller, 1998; Olypher et al., 2002; Jackson and Redish, 2007). PCP increased overdispersion, which was greatest after the moderate dose (F_2,169_ = 3.51, p = 0.032, ANOVA; Fig. 4F). Given that reduced overdispersion correlates with singularly-focused spatial attention (Fenton et al., 2010), the overdispersion increase raises the possibility that PCP interferes with attention, a crucial component of cognitive control. Increased overdispersion also indicates that despite preserved place fields, moment-to-moment, the ensemble representation of space was excessively variable, suggesting degraded position representation in ensemble discharge.

### Impaired position representation under PCP

We then assessed whether spatial information in place cell discharge is degraded following the moderate dose of PCP using a Bayesian algorithm to decode the rat’s position from the momentary (300 ms) discharge of single unit ensembles (Zhang et al., 1998). Only ensembles with at least 5 place cells (avg: 8.4 ± 2.3 cells) were examined. The average posterior probabilities appeared elevated over a larger area after the moderate PCP dose (Fig. 6A), suggesting greater inaccuracy. This was quantified by computing the decoding error as the distance between the decoded and observed locations (see example distributions Fig. 6B top) then comparing the average decoding error before and after the various PCP injections (Fig. 6B bottom). The average decoding error in the post-injections recordings were Control = 20.6 ± 3.6 cm, Low = 21.6 ± 3.2 cm and Moderate = 26.2 ± 6.8 cm PCP doses, which was significantly increased compared to the respective pre-injection error after the moderate PCP dose (F_2,12_ = 4.2, p = 0.041, ANOVA; Dunnett’s test: Control = Low < Moderate). The confidence in the position decoding was also decreased by the moderate PCP dose, as estimated by the posterior confidence area, the region with elevated probability (Fig. 6C top). Compared to the respective pre-injection recordings the effect of PCP dose was significant (Fig. 5C bottom; F_2,12_ = 4.3, p = 0.039, ANOVA; Dunnett’s test: Control = Low < Moderate).

### CA1 discharge changes in the LFP theta phase preference under PCP

Next we examined the relationships between the LFP and place cell discharge. Analysis of the frequency-specific phase at which cells discharge, revealed cell-specific patterns of firing that were observed by plotting the probability of discharge within the temporal space defined by the phase and frequency of oscillations in the ongoing LFP (Fig. 7A). By inspection, the CA1 discharge is most strongly phase-organized by theta-frequency oscillations, but the specific phase relationship varies across cells. Gamma-phase organization is less prominent (Fig. 7A). PCP changed this temporal organization of discharge (F_2,160_ = 3.1; p = 0.048, ANOVA; Fig. 7B), specifically at theta frequencies. The 2-way ANOVA comparing the phase-frequency-discharge probability plots before and after PCP administration in the theta, slow, and medium gamma ranges confirmed significant effects of PCP dose (F_2,482_ = 9.2; p = 1.20×10^−4^, ANOVA) and the frequency band (F_2,482_ = 3.3; p = 0.038, ANOVA) but not their interaction (F_4,482_ = 0.4; p = 0.809, ANOVA). Post-hoc tests confirmed that phase-frequency discharge probability changes from baseline were only caused by the moderate dose of PCP (Fig. 7B) and the effect was selective for theta band oscillations (Fig. 7C).

Next we investigated if PCP caused a uniform change in the phase-frequency discharge probability of the CA1 cells or whether PCP discoordinated the relationships that the network of place cells has to the LFP. This was estimated by computing the set of pairwise relationships between place cell firing and oscillations in the LFP. As shown in Figure 7D, a cell pair’s the difference in the phase-frequency discharge probability profiles for a cell pair was similar before and after PCP injections (Pearson correlation coefficient r’s range -0.303 to 0.889, d.f.’s = 1998, p’s range 0.003 to 8.20×10^−680^; Fig. 7D), but the change was greatest after the moderate dose of PCP, specifically in the theta band (Fig. 7E). The 2-way ANOVA confirmed a significant dose and band interaction (F_4,1787_ = 5.7; p =1.47×10^−4^, ANOVA) as well as significant effects of dose (F_2,1787_ = 16.7; p = 6.52×10^−8^, ANOVA) and band (F_2,1787_ = 88.4; p = 2.24×^-37^, ANOVA). In addition to discoordinating oscillations in the LFP, these findings confirm that the moderate dose of PCP discoordinates the timing of CA1 place cell discharge with respect to the phase of theta oscillations.

### Discoordinated action potential discharge under PCP

Next we examined if PCP discoordinates neural discharge amongst cells on the theta timescale of 250 ms. Because the rats were typically moving 5.87 ± 0.21 cm/s and moved faster during the modest dose of PCP (10.05 ± 1.28 cm/s), during 250 ms they are mostly detected within a single 2.5 cm × 2.5 cm location, which is the spatial resolution of the firing rate maps we computed. This analysis measures neural coordination that can only weakly be affected by the increased movement of PCP-treated rats; they rarely changed the ~6 cm^2^ analysis location during the 250 ms analysis interval (Olypher et al., 2003). Accordingly, compared to analyses that depend on averaging across longer time scales and position, this short-time scale, location-independent analysis is relatively insensitive to hyperlocomotion. The Kendall correlations of all simultaneously recorded cell pairs of place cells (Fig. 8) shows that the discharge of some cell pairs is positively correlated on the time scale of a few hundred milliseconds, whereas the discharge of other pairs is uncorrelated or negatively-correlated (Olypher et al., 2006; Kelemen and Fenton, 2010; Kelemen and Fenton, 2012).

We next examined whether PCP alters the coupling of discharge within the network of CA1 place cells. This was motivated by theoretical and empirical evidence that cognitive control is impaired by selectively increasing initially low correlations in ensemble action potential discharge (Wesierska et al., 2005; Olypher et al., 2006; Murray et al., 2014). MK-801, another NMDAR psychotomimetic, disrupts correlated firing in the prelimbic region of medial prefrontal cortex (Molina et al., 2014) as well as hippocampal CA1 ensembles recorded under urethane anesthesia (Szczurowska et al., 2017). Ketamine, another NMDAR psychotomimetic also increases interregional co-fluctuations of fMRI signals (Driesen et al., 2013). The stability of neural coordination was estimated by PCo, the Pearson correlation (r) comparing cell-pair coordination (measured as Kendall’s τ) between two 10-min epochs (Neymotin et al., 2017). Before injection, the coordination of cell pairs was stable across the three 10-min epochs (Dose: F_2,615_ = 1.08, p = 0.340, ANOVA; Epoch: F_1,615_ = 0.44, p = 0.507, ANOVA; Interaction: F_2,615_ = 1.40, p = 0.247, ANOVA; Fig. 8A). The moderate dose of PCP decreased the stability of the coordination but the control and low doses did not change PCo (Dose: F_2,642_ = 5.00, p = 0.007, ANOVA; Epoch: F_1,642_ = 2.05, p = 0.153, ANOVA; Interaction: F_2,642_ = 3.09, p = 0.046, ANOVA; significant post-hoc comparisons: Moderate pre-injection < Moderate post-injection; Control post-injection < Moderate post-injection; Fig. 8B). The coordination of cell pairs was stable after injection but higher for the moderate dose (Dose: F_2,642_ = 13.14, p = 2.55×10^−5^, ANOVA; Epoch: F_1,642_ = 0.13; p = 0.719, ANOVA; Interaction: F_2,642_ = 0.26, p = 0.771, ANOVA; Fig. 8C). To analyze the stability of coordination across the different pre- and postinjection comparisons, the change of coordination between the early and late epochs was first normalized by the standard deviation of the pre-injection **τ** values (i.e. [**τ**_late_ - **τ**_early_]/s.d.[**τ**_early_]). The moderate dose of PCP increased the coupling of cell pair discharge but the control and low doses did not cause changes (Dose: F_2,963_ = 2.32, p = 0.099, ANOVA; Comparison (Pre-Pre, PrePost, Post-Post): F_2,963_ = 1.04, p = 0.354, ANOVA; Interaction: F_4,963_ = 4.67, p = 9.76×10^−4^, ANOVA; Post-hoc comparisons: the moderate PCP pre-post change is significantly different from all other changes, p’s < 0.03; Fig. 8D).

Despite the robust findings of neural discoordination from analyses on sub-second timescales we further investigated whether the findings arise from hyperlocomotion expressed on the seconds time scale resulting in the traversal of more firing fields. We first confirmed that despite a dose- dependent effect on hyperlocomotion, only the moderate dose of PCP administration changed the spike train correlations amongst pairs of simultaneously recorded complex-spike cells, regardless of spatial firing quality and how close or far apart the firing fields were located (Fig. 9A; F_2,660_ = 5.8, p = 0.003, ANOVA). We also recorded 11 CA1 complex-spike cells in one additional rat that was restricted to a small 24-cm diameter enclosure, which minimized locomotor movements and therefore should have attenuated any effects that were consequences of PCP-induced hyperlocomotion (Fig. 9B). The set of 53 cell-pair correlations in this rat did not significantly change in the subset of cell pairs that were independent or positively correlated in the baseline recordings but the moderate dose of PCP increased correlated discharge amongst cell pairs that were negatively correlated prior to drug administration (53 cell pairs, 2 pairs were extreme in baseline and excluded from analysis; effects of: PCP treatment F_1,102_ = 4.80, p = 0.031, ANOVA; baseline cell pair category F_2,102_ = 11.64, p = 2.80×10^−4^, ANOVA; interaction F_2,102_ = 6.24, p = 2.78×10^−3^, ANOVA). After PCP treatment 2/3 of negatively correlated cell pairs increased by more than two s.d., whereas only 3/53 cell pairs changed in the independent and positively correlated categories (test of proportions z = 4.5, p = 1.60×10^−5^). This pattern of increased and maintained correlations has been observed under urethane anesthesia after disinhibiting the hippocampus by injecting tetrodotoxin into the contralateral hippocampus (Olypher et al., 2006), a manipulation in awake rats that impairs learning, consolidation, and recollection of the Room+Arena- active place avoidance task variant with significant cognitive control demand but not the Room+ task variant with reduced cognitive control demand (Wesierska et al., 2005), resembling what we observed here with PCP (Fig. 2).

Finally, to completely rule out that these PCP-induced changes in neural coordination were a consequence of the PCP effect on locomotion, we investigated whether PCP administration also causes neural discoordination under urethane-anesthesia (Fig. 9C). Similar to the findings in the awake rat, few cell pairs had initially negative correlations, and after PCP their correlated discharge increased (8/10 by > 2 s.d.) together with the independent cell pairs (15/23 by > 2 s.d.). In contrast, there was no effect on cell pairs with initially positive correlations (4/47 changed by > 2 standard deviations; test of proportions z’s > 8.1, p = 2.26×10^−15^). The main effects as well as the interaction between PCP treatment and baseline cell pair category were significant (effect of PCP: F_2,147_ = 6.22, p = 0.014; effect of baseline category: F_2,147_ = 11.18, p = 3.021×10^−5^; interaction: F_2,147_ = 4.23, p = 0.016, ANOVA). We also made these recordings from mPFC, where in contrast to hippocampus, almost all cell pairs had significantly positively correlated responses (113/127; Fig. 9D). After the moderate dose of PCP, 3/36 of the positively correlated cell pairs increased by > 2 s.d., whereas all cell pairs increased their correlation if they were independent (2/2) or negatively (2/2) correlated before PCP. We observed positively, negatively and independently coupled hippocampus-mPFC cell pairs (n = 250; Fig. 9E). PCP selectively increased the correlation between the initially independent and negatively coupled cell pairs (effect of PCP F_1,244_ = 8.77, p = 3.37×10^−3^, ANOVA; effect of baseline category F_2,244_ = 8.19, p = 3.62×10^−4^, ANOVA). Thus, in both the dorsal hippocampal and mPFC networks, PCP preferentially increased the coactivation of cell pairs that were weakly coactive before PCP. Together, these data strongly support the view that the moderate dose of PCP changes neural coordination independently of behavior.

Since the increases in weak pair-wise correlations is pathognomonic of strong changes in neural network states (Schneidman et al., 2006), we next examined the discharge pattern across the entire place cell ensemble as a function of recording time before and after PCP administration. In an example recording (Fig. 10A), prior to PCP administration, the ensemble activity pattern during each minute was similar to the pattern at each other minute (Fig. 10B, C), which was quantified by high, significant Pearson’s correlation coefficients (r ~ 0.7) of the ensemble activity vectors. The moderate dose of PCP radically changed this stationarity in the temporal structure of ensemble activity. By the first post-PCP minute, the ensemble pattern was uncorrelated (Pearson’s correlation coefficient r ~ 0.0) to the pre-injection pattern. The post-PCP ensemble pattern was at least as stationary as the pre-injection pattern for over 20 minutes. From about 25 minutes after PCP, the ensemble activity switched between the pre- and post-injection patterns at intervals of several minutes (Fig. 10B,C). The moderate dose of PCP consistently induced an aberrant pattern of ensemble discharge that did not correspond to the established representation. This abnormality was not observed following the control or the low doses of PCP (Fig. 10D-G; F_2,18_ = 11.93, p = 5.03×10^−4^, ANOVA). While this seconds-scale discoordination of ensemble neural discharge is predicted by the increased pair-wise correlations at 250 ms resolution, it could not be explained by the PCP-induced changes of spatial behavior (Fig. 11). Thus, the moderate dose of PCP discoordinated the temporal relationships between pairs and ensembles of cells, but the location-specific firing of individual place cells remained relatively undisturbed (Fig. 4).

Furthermore, the Fig. 11A correlation matrix suggests that CA1 activity is multistable and that PCP may simply change the likelihood of observing different ensemble activity states, rather than inducing a *de novo* activity pattern. In this example, during the baseline, prior to PCP, the ensemble activity was transiently more like the PCP activity than the baseline activity. Such occasions are visible as moments of low correlation (yellow stripes) with the first 30 minutes of pre-PCP activity vectors and as moments of high correlation (red stripes) with the 30 minutes of post-PCP activity vectors. Reminiscent of the preplay phenomenon (Dragoi and Tonegawa, 2011, 2013), this observation suggests that both the pre-PCP and post-PCP neural activity had been encoded in CA1, and that experiencing PCP merely increased the probability of expressing the pre-established post-PCP pattern.

## DISCUSSION

### Main findings

We find that a moderate dose of PCP that impairs spatial cognitive behavior that requires cognitive control (Fig. 2), and discoordinates the timing between distinct hippocampal CA1 signals with minimal disruption of place fields (Figs. 4,5). The Fig. 4 and 5 differences in the quantitative measures of firing field quality and stability illustrate the impact of position sampling on these measures, as has been reported (Muller et al., 1987; Maurer et al., 2005). This discoordination was measured in multiple ways, 1) as an aberrant dominance of neocortical input-associated medium gamma oscillations over CA3 input-associated slow gamma oscillations (Fig. 3); 2) as a disorganization of spiking by theta oscillations (Fig. 7); 3) as a tendency of place cells that did not discharge together to increase sub-second coactive firing, which manifested as strengthened and unfamiliar hippocampal network states during behavior in a familiar environment (Figs. 8,9,10). This was also observed in hippocampus-mPFC investigations of neural coordination (Fig. 9). Despite the neural discoordination, PCP had minimal effects on the time-averaged spatial firing properties of individual place cells. This spatially accurate firing became more temporally unreliable under PCP on the time scale of the few seconds of crossing a firing field (Fig. 4F). Consistent with this temporal unreliability, decoding location from ensemble discharge on the 300 ms time scale was worse after the moderate dose of PCP (Fig. 6). These studies demonstrate that PCP-induced cognitive impairment and neural discoordination can be independent of psychotomimetic-induced hyperlocomotion, unlike most prior studies. While the hyperlocomotion was general, the effects of PCP on cognitive behavior and neural activity were task specific (Fig. 2D), frequency-band specific (Figs. 2H, 3B-D), and neural-coordination category specific (Figs. 7C-E, 8). Furthermore, some forms of neural discoordination could also be demonstrated under anesthesia (Fig. 9C-E) and at sub-second time scales during which the spatial behavior was effectively constant during the window of analysis (Figs. 8,9).

These findings suggest that memory storage itself was not disturbed by PCP and demonstrate stable hippocampus place fields during impaired hippocampus-dependent spatial cognition. The findings highlight that normal, stable place fields do not predict normal spatial competence. We observed instead that the dose of PCP that induces neural discoordination also impairs a familiar place avoidance task, but only if the demand for cognitive control is high (Fig. 2). This indicates that PCP-induced deficits are in the use of established spatial memories, rather than in place learning or information storage *per se* (Bannerman et al., 2012). Consistent with unimpaired Room+ place avoidance under PCP, the rat’s spatial competence recovers once the acute effects of PCP wear off. Furthermore, directly infusing PCP into dorsal hippocampus impairs familiar place avoidance and the magnitude of the impairment is indexed by the aberrant increase of medium gamma oscillations (Fig. 2H), demonstrating the action of PCP in the hippocampus is sufficient to cause these cognitive consequences but not the sensorimotor disturbances that accompany systemic PCP administration (Fig. 2C).

### Electrophysiological effects of PCP

The impairing dose of PCP enhanced the theta-phase modulation of medium gamma without changing the modulation of slow gamma. Medium gamma in CA1 is associated with neocortical activity and slow gamma with memory-related CA3 input (Colgin et al., 2009; Bieri et al., 2014; Schomburg et al., 2014). Accordingly, the present findings suggest that PCP affects the likelihood of processing in the separate neocortical and hippocampal information streams. Such a PCP-induced imbalance may account for the inability to resolve conflicts between relevant and irrelevant spatial information in the Room+Arena- task variant, even when the conditioned avoidance is familiar. The increase in gamma power we observed replicates prior work that PCP (Ma and Leung, 2000), and other NMDAR antagonists increase gamma oscillations (Ehrlichman et al., 2009; Hong et al., 2010; Lazarewicz et al., 2010), as well as reports of altered theta- medium gamma cross-frequency coupling (Caixeta et al., 2013) (Fig. 3). The present observations extend these findings to destabilized cell-specific theta phase discharge probabilities (Fig. 7), and increased discharge coupling amongst initially weakly coupled cell pairs (Figs. 8,9) to generate unfamiliar and strong network states (Fig. 10). Taken together, the major effect of PCP is to discoordinate neural activity, an effect that has been reported under anesthesia with MK-801 a related drug, in hippocampus (Szczurowska et al., 2017) as well as mPFC (Molina et al., 2014). Preserved place cell spatial firing in familiar conditions after PCP (Fig. 4) is consistent with prior work using the competitive NMDAR antagonist CPP (Kentros et al., 1998; Ekstrom et al., 2001). Yet the temporal organization of discharge amongst groups of CA1 cells was changed on multiple time scales ranging from 100’s of milliseconds (Figs. 7, 8, 9) to minutes (Fig. 10). Preserved place cell firing after PCP is consistent with spared spatial memory and navigation after NMDAR antagonists (Morris et al., 1986; Bannerman et al., 1995; Saucier and Cain, 1995), while the discoordination of neural discharge predicted the PCP– induced impairment of cognitive control.

Finally, an unexpected observation suggested that PCP could merely increase the probability of expressing latent but unusual ensemble patterns of activity (Fig. 11A). This is intriguing because PCP induces hallucination (Cohen et al., 1962; Abi-Saab et al., 1998). But how might hallucination manifest in a rat? A mechanistic hypothesis for a neural correlate inspired by the Fig. 11A observation is that such unusual ensemble activity patterns are an alternative expression of the otherwise normal activity of individual place cells. While substantial further work is required to investigate such a possibility and others, the place cell research paradigm may offer a means to rigorously investigate the potential relationship between such PCP-induced dynamics and hallucination (Hoffman, 1997; Hoffman and McGlashan, 2001). These electrophysiological findings highlight what has been called “the problem of the missing middle.” Relatively little is known about how the well-characterized molecular details of PCP’s actions alter mesoscale neural network processes to produce the also relatively well-characterized macroscale observables of behavior (Laughlin et al., 2000).

### Novel potential motivation and utility for PCP and psychomimetic-based investigations

Acute PCP intoxication has been the most extensively used experimental preparation for antipsychotic drug development that focused on treating the positive symptoms of psychosis. As the drug development focus shifted from attenuating psychosis to cognition promoting treatments, interest in the PCP model has waned, also due to the availability of more selective antagonist compounds and genetic models of potential etiological contributors to schizophrenia. Given the current interest to develop drugs that target cognitive symptoms in schizophrenia, the present demonstration of a set of PCP-induced cognition related deficits indicates that the use of the PCP model could also be extended to the search for a procognitive antipsychotic (Moghaddam and Krystal, 2012). Indeed, recordings in prefrontal cortex under PCP and related NMDAR antagonist psychotomimetics indicate the drugs discoordinate prefrontal network discharge and gamma oscillations (Wood et al., 2012), observations that the present findings extend to hippocampus and demonstrate with direct evidence of discoordinated neural representations of place and spatial information in the awake rat, as well as under anesthesia. As such, the relationship between PCP induced discoordination in dorsal and ventral hippocampus and prefrontal cortex merits further study (Jodo et al., 2005; Jodo, 2013).

It is important to consider the dissociation between the novel PCP-induced cognitive control deficit (Fig. 2D) and the long-established hyperactivity that is thought to model the positive symptoms (Fig. 2C). Infusing PCP directly into the hippocampus impaired cognitive control measured by place avoidance (Fig. 2G) without causing hyperactivity (Fig. 2F), so long as the drug sufficiently increased medium gamma oscillations (Fig. 2H). The dissociation could help understand why treatment strategies that were developed by targeting hyperactivity and other sensorimotor abnormalities turned out to be inadequate for the cognitive symptoms (Carter and Barch, 2007). The present work also provides compelling evidence that although NMDAR antagonism spares established memory, it can devastate the judicious use of information when sources of cognitive interference abound such as during Room+Arena- place avoidance or initial place learning in the water maze (Bannerman et al., 1995; Saucier and Cain, 1995; Bannerman et al., 2006). These findings suggest it may be fruitful to investigate cognitive control, in addition to learning and memory *per se*, in research directed at schizophrenia as well as other fields. While these behavioral findings on their own suggest that acute PCP intoxication could with the right assays be exploited to model the core cognitive deficits in schizophrenia, the utility of the model lies in the theoretical importance of the findings. The electrophysiological findings provide unambiguous evidence that PCP discoordinates electrical neural activity between neurons (Figs. 6–11) with minimal consequences on the response characteristics of individual cells (Tables 1, 2; Figs. 4,5) as predicted by the discoordination hypothesis. The findings that doses of PCP that impair cognitive behavior also discoordinate neural activity suggest that normalizing neural disccordination could be a procognitive therapeutic target. Although this may be accomplished by correcting a proximal cause of the dysfunction, procognitive effects can also be achieved by correcting the discoordination itself, using pharmacological and/or neuromodulation treatment strategies to restore excitation-inhibition coordination, or even by harnessing the plasticity that can accompany appropriate cognitive experience (Lee et al., 2012; Fenton, 2015).

## AUTHOR CONTRIBUTIONS

Data were collected by H-YK, JK and EP analyzed by H-YK, EP, DD, EK and AAF; AAF supervised research; H-YK and AAF designed research; AAF wrote the paper and all authors contributed to editing and figures.

## ACKNOWLEDGEMENTS

Supported by NIMH grants R21MH082417 and R01MH084038. We thank Steven E. Fox for comments on the manuscript.

